# Formins and Arp2/3 Reciprocally Regulate Contact Guidance on Aligned Collagen Fibrils

**DOI:** 10.1101/2025.01.03.631234

**Authors:** Azarnoosh Foroozandehfar, Shayan Tohidi, Shafayet Ahmed Siddiqui, Fred Rogers Namanda, Michael Forrester, Eric Cochran, Ian C. Schneider

## Abstract

Directed cell migration is essential in many biological processes and is driven by a variety of directional cues, including aligned fibrils in the extracellular matrix (ECM), a phenomenon known as contact guidance. How different cells respond to aligned fibrils and how internal regulators like formins and Arp2/3 control contact guidance is unknown. In this study, a unique system to assemble aligned collagen fibrils on mica and to transfer them onto controllable substrates is used to probe contact guidance. This fibril alignment system reveals that cytoskeletal regulation through myosin contractility and not receptor expression drives contact guidance ability. Highly contractile cells exhibit high-fidelity contact guidance, weakly contractile cells ignore cues and moderately contractile cells use a mixture of both parallel and perpendicular migration strategies on aligned collagen fibrils. In addition to myosin contractility, formins and Arp2/3 control contact guidance in a reciprocal manner across a variety of cell types. Formins, mediators of linear F-actin structures, enhance contact guidance and Arp2/3, a mediator of branched F-actin structures, diminishes contact guidance. This controlled materials system reveals the importance of both myosin-mediated contractility as well as the antagonistic action of formins and Arp2/3 on contact guidance, providing potential targets to tune contact guidance.

## 1. Introduction

Contact guidance refers to the directed migration of cells along aligned fibrils present in the extracellular matrix (ECM). Studying contact guidance is particularly important because contact guidance controls embryonic development ^[1,2]^ and wound healing ^[3,4]^. Contact guidance also impacts pathological conditions, directing migration in fibrosis and during cancer metastasis.^[5,6]^ Contact guidance can be recreated *in vitro* using various methods, including electrospinning,^[7]^ which produces synthetic aligned fibrils; microcontact printing,^[8,9]^ which produces printed lines of ECM and grooves or ridges ^[10]^ that guide cellular movement topographically. While these systems recapitulate certain geometric and chemical features, they lack ultrastructural signatures or the ability to present cryptic ligand binding cites^[11]^ that are found on collagen fibrils assembled *in vivo*. Approaches exist to make aligned collagen fibrils, including magnetic field alignment,^[12]^ mechanical rotation,^[13]^ flow-based alignment ^[14,15]^ or cell-based alignment.^[16–18]^ However, these 3D systems can be difficult to control and present challenges in imaging. Our lab has used a 2D self-assembly approach to make aligned collagen fibrils through epitaxial growth on mica, resulting in fibrils closely resembling those in tissues. D-banding, a ultrastructural feature that is seen on these substrates, is critical for contact guidance.^[19]^ We have developed methods to finely tune the structure of collagen fibrils on mica.^[20]^ Cell-type specific differences in contact guidance that we observe in this 2D self-assembly system aren’t visible using other 2D contact guidance systems, but faithfully replicate contact guidance behavior seen in 3D collagen gels.^[13,21]^ However, collagen fibrils assembled on mica tend to delaminate when exposed to cells under certain conditions.^[22]^ Consequently, while epitaxial growth of aligned collagen fibrils on mica affords a powerful approach to study contact guidance that retains ultrastructural characteristics of the collagen fibrils and mimics behavior seen in less tunable 3D contact guidance environments challenges associated with collagen fibril delamination remain.

Even though contact guidance cues can be presented to cells, cells may perceive that directional information differently. Differing roles within tissues likely results in different contact guidance behavior amongst cell types. For instance, fibroblasts must remodel the ECM and generate contractile forces to provide mechanical stability of the wound, resulting in transient alignment of the ECM within the wound bed that enhances cell infiltration, including additional fibroblasts via contact guidance.^[3,23]^ Over the long term, this alignment is destroyed once mechanical integrity and directed cell migration are no longer needed.^[24]^ Epidermal keratinocytes on the other hand facilitate re-epithelialization outward in and may need to be less sensitive to fibril alignment, given that wounds frequently re-epithelialize uniformly from all directions.^[3]^ Finally, immune cells need to randomly surveil the wound microenvironment with no bias in migration in response to aligned collagen fibrils, requiring them to ignore contact guidance cues. Indeed, ECM fibril alignment seems not to enhance T cell infiltration into the tumor ^[25]^ and T cell contact guidance is low compared to other cell types.^[26]^ It is not fully understood how different cells engage in contact guidance and what intracellular pathways are overlapping and what pathways are distinct.

While early contact guidance work did not delineate between differences among cells of different type, more recent work suggests that there are some differences in how cells sense and respond to contact guidance cues.^[27–30]^ Different qualitative modes of migration have been described that have allowed for a coarse-grain understanding of cell migration. Mesenchymal cells adopt a spindle shape, showing strong adhesion and many stress fibrils, usually oriented parallel to the direction of migration. Epithelial cells adopt a half-moon shape with a broad lamellipodium, intermediate adhesive strength, and stress fibrils both parallel and perpendicular to the direction of migration. Amoeboid cells are rounded with low adhesion and few stress fibers as well as a cortical contractile and protrusive F-actin cytoskeleton. In addition, there are cells that can irreversibly or dynamically switch between epithelial and mesenchymal migration modes as well as those that switch between amoeboid and mesenchymal modes of migration. ^[31,32]^ Mesenchymal cells showed higher contact guidance than those undergoing epithelial-to-mesenchymal transition (EMT).^[29,33]^ Ameboid cells showed random alignment, while mesenchymal cells showed directed migration in both 2D and 3D systems.^[13,21]^ While the contact guidance of different cells has been assessed, nearly all this work has been completed on substrates that may obscure cell type differences *in vivo.*^[21]^ Furthermore, what intracellular processes are overlapping and what are distinct between different cell types is not known. In particular, how epithelial cells respond to contact guidance cues and whether this response mimics what is seen in mesenchymal cells is not understood.

Differences in contact guidance across different cells could be attributed to differences in surface receptors, but in instances where cells spread and migrate on a particular substrate, likely differences in contact guidance are due to cytoskeletal regulation. While increased actomyosin contractility has been suggested to enhance contact guidance, how other regulators of the F-actin cytoskeleton control contact guidance is not known. Cells polymerize both linear and branched F-actin networks that allow the cell to migrate directionally. Formins, promote the formation of linear actin filaments, which are essential for creating stress fibrils and contractile forces that drive cell body movement and stabilize cell adhesion.

Arp2/3 on the other hand is responsible for forming branched actin networks, which are crucial for generating the pushing forces needed for forming lamellipodia and protrusions at the leading edge of migrating cells. Since both formins and the Arp2/3 complex polymerize F-actin from the same G-actin pool, others have suggested competition between these pathways might result in reciprocal control over cytoskeletal functions.^[34]^ We hypothesize that this competition results in reciprocal control over contact guidance by formins and the Arp2/3 complex. Since formins build linear F-actin networks that are found during contact guidance, it is reasonable to hypothesize that formins positively regulate contact guidance. The role of the Arp2/3 complex in contact guidance is unclear due to conflicting results in literature.^[35–37]^ This work has been carried out in contact guidance systems that do not present contact guidance cues as fibrils, potentially masking the true role of Arp2/3. Consequently, it is unknown whether formins and Arp2/3 reciprocally regulate contact guidance and whether this is a shared mechanism across cells with different modes of migration.

In this paper, we optimize a fibril transfer technique to move native collagen fibrils assembled on mica onto other substrates, blocking collagen delamination by highly contractile cells. The contact guidance response of different cells with distinct migration modes is explored. We systematically assess whether receptor expression or cytoskeletal regulation better predict contact guidance fidelity. We probe the role of formin and Arp2/3-mediated F-actin polymerization in determining contact guidance fidelity to test the hypothesis that contact guidance is reciprocally controlled by formins and Arp2/3. Finally, we characterize a newly discovered epithelial migration mode on aligned collagen fibrils. This mode of migration involves switching between parallel and perpendicular migration with respect to aligned collagen fibrils. We explore the balance of formin-mediated linear F-actin polymerization and Arp2/3- mediated branched F-actin polymerization on parallel and perpendicular migration of epithelial cells.

## 2. Results

### 2.1 Self-assembled collagen fibrils transferred to glass resist collagen fibril delamination and induce robust fibroblast alignment

Collagen fibrils can be epitaxially assembled on mica via electrostatic interactions between the collagen fibrils and the mica. Under specific buffer conditions which include collagen concentration and adsorption time, well-aligned collagen fibrils can be formed and imaged using AFM. *In vivo* characteristics like D-banding is observed (**Figure 1A**). Human foreskin fibroblasts (HFFs) initially align along the collagen fibrils, however, over time they contracted and delaminated the collagen fibrils shown as large aggregates at 4 hours (**Figure 1A**). The kinetics of delamination were measured as the fraction of initial HFFs remaining on the substrate over time (**Figure 1B**). A statistically significant decrease in the fraction of HFFs remaining occurred after 2 hours. This creates a significant problem with using these substrates for highly contractile cells. Consequently, we assessed whether delamination occurred on other substrates. We adsorbed collagen trimers (t) to mica using low pH, assembled aligned collagen fibrils (af) on mica, created multi-directional collagen fibrils (mdf) by using higher collagen concentrations and transferred aligned collagen fibrils (tf) to functionalized glass. We previously explored a technique to transfer these fibrils onto other substrates,^[38]^ but reliability in the transfer approach and proper quantification of the transfer efficiency was not completed. AFM images verified that the solution conditions affect the collagen structure and that the optimized collagen fiber transfer approach faithfully transferred aligned collagen fibrils (**Figure 1C and Supplementary Figure 1**). Cell alignment was quantified using a directionality index. Directionality index values of 1 indicate high alignment with collagen and high-fidelity contact guidance and values of 0 indicate low alignment with collagen and low contact guidance fidelity. HFFs were not well spread or aligned on collagen trimers (t) (**Figure 1D&E**). On aligned collagen fibrils assembled on mica, HFFs were aligned well initially (1 h), but eventually detached due to collagen delamination (4 h), leaving both large clusters as well as a small number of single cells that were not well aligned (**Figure 1D&E**). Multi-directional collagen fibrils induced better spreading than the collagen trimers, but the cell appeared confused and directional alignment was poor.

**Figure 1.**
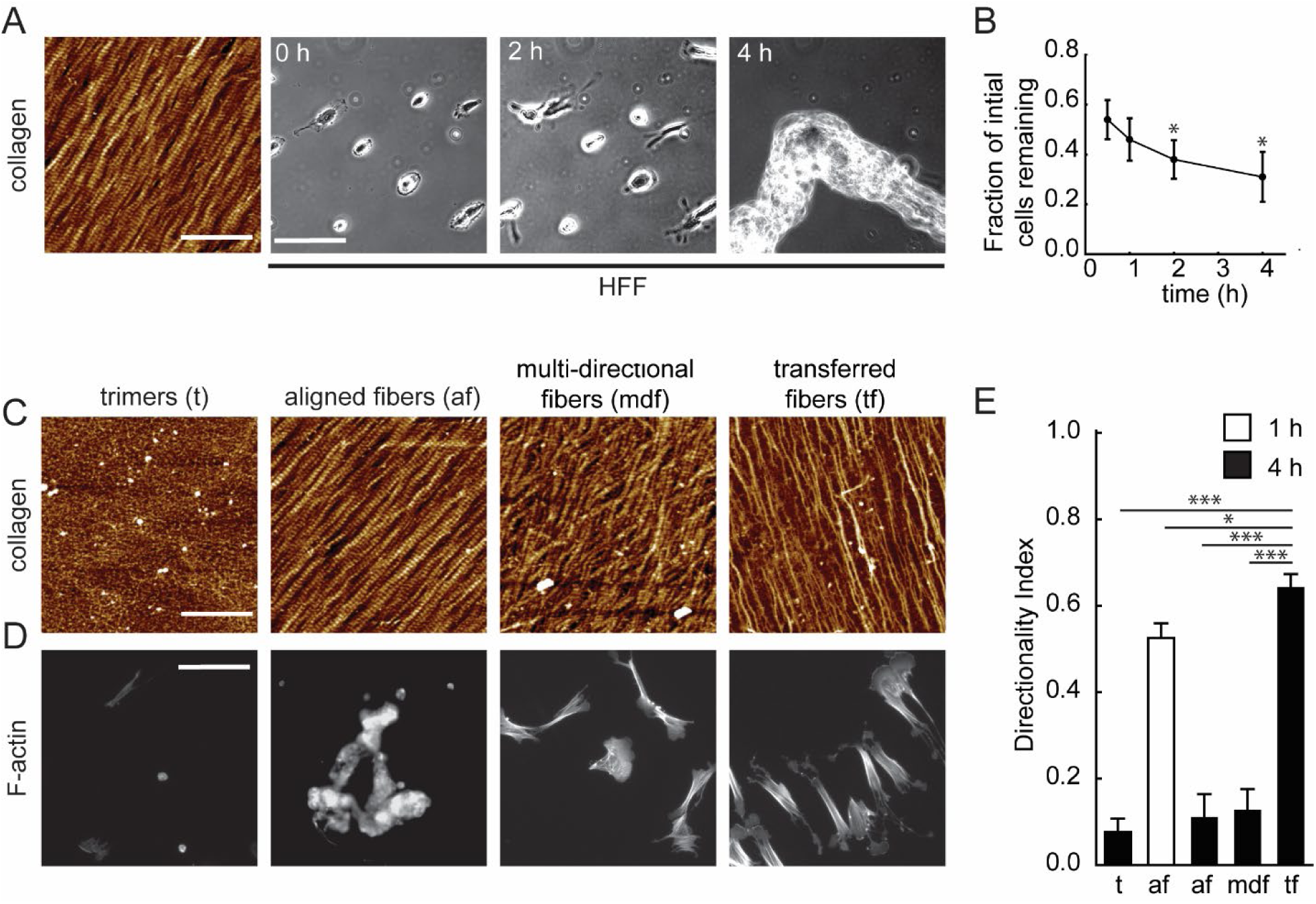
Contractile fibroblasts respond differently on different collagen substrates. (**A**) AFM image of collagen assembled on mica. AFM scale bar is 1 μm. Live-cell phase contrast microscopy images of HFFs on mica at 0, 2 and 4 h. Phase contrast image scale bar is 150 μm. (**B**) Fraction of initial cells remaining on the mica surface over time as assessed after fixation. *N_samples_* ≥ 4, *N_cells_* ≥ 304 and *N_images_* ≥ 20. Statistical significance was assessed by comparing to the 0.5-hour time point using a one-way ANOVA (single factor), where * denotes *p* ≤ 0.05. (**C**) Various collagen patterns assembled on mica under different conditions. Trimers (t): citric acid buffer (pH 4.3) with 200 mM KCl, 10 µg/ml collagen and a 12 hours incubation. Aligned fibrils (af): 50 mM Tris buffer (pH 9.2) with 200 mM KCl, 10 µg/ml collagen and a 12 hours incubation. Multi-directional fibrils (mdf): 50 mM Tris buffer (pH 9.2) with 200 mM KCl, 100 µg/ml collagen and a 12 hours incubation. Transferred fibrils (tf): 50 mM Tris buffer (pH 9.2) with 200 mM KCl, 10 µg/ml collagen, a 12 hour incubation and transferred onto functionalized glass. Scale bar is 1 μm. (**D**) Phalloidin staining showing F-actin structure and HFF morphology on collagen fibrils under the same conditions as in **C**. Scale bar is 150 μm. (**E**) Directionality index of HFFs on different collagen patterns and after different incubation times on the collagen fibrils. *N_samples_* ≥ 3 and *N_cells_* ≥ 304. Statistical significance was assessed by comparing to collagen fibers transferred to glass, where * denotes *p* ≤ 0.05, ** denotes *p* **≤** 0.01 and *** denotes *p* ≤ 0.001. Error bars are 95% confidence intervals.

Transferred collagen fibrils resulted in robust spreading and alignment of HFFs due to the chemical linkage of the collagen fibrils to the glass surface with no delamination of the collagen fibrils after 4 hours (**Figure 1E**). Consequently, HFFs, mesenchymal cells that are known to be highly contractile cells engage in robust contact guidance on aligned collagen fibrils, but not on other non-aligned collagen substrates. Furthermore, collagen delamination and deadhesion of cells is blocked when fibers are transferred to glass, allowing for long-term contact guidance studies.

### 2.2 Regulation of F-actin cytoskeleton and not collagen receptor expression explain differences in contact guidance among cell types

Given that HFFs robustly spread and orient on aligned collagen fibrils, but only when collagen fibril delamination was blocked by transferring collagen fibrils to glass, we examined whether this behavior was observed in other cell lines with different modes of migration. Previously, we showed that carcinoma cells that adopt a mesenchymal mode of migration (MDA-MB-231) tended to robustly orient and directionally migrate on aligned collagen fibrils, whereas carcinoma cells that adopt an amoeboid mode of migration (MTLn3) did not.^[20,21]^ Along with HFF and MDA-MB-231 cells, we examined WM-266-4, HaCaT, and THP-1 cells. WM-266-4 cells are a mesenchymal to amoeboid mode-switching cell line.^[32]^ HaCaTs migrate as epithelial cells. THP-1 cells are a model naive immune cell that moves with an amoeboid migration mode. We examined the orientation of individual cells across different cell lines on collagen fibrils assembled on mica as well as those transferred to glass (**Figure 2A**). The delamination of collagen fibrils assembled on mica in response to HFF (**Figure 1**) is blocked when collagen fibrils are transferred to glass. This suggests that the perceived stiffness is lower on fibrils adsorbed to mica compared to those transferred to glass, controlling the cell response to aligned collagen fibrils. HFF and MDA-MB-231 cells adopted extended, large area morphologies with long stress fibers in the direction of the long axis of the cell, typical of mesenchymal cells. However, HFF cells had a wider body and more pronounced F-actin stress fibrils within and paxillin focal adhesions underneath the cell body (**Figure 2B**). WM-266-4 and HaCaT cells adopted an epithelial cell morphology with numerous different types of F-actin fibrils including longitudinal stress fibers and transverse arcs that terminated in large focal adhesions (**Figure 2B**). Finally, THP-1 cells were not well spread, did not include well-defined F-actin fibers and only had small focal adhesions (**Figure 2B**). HFF cells spread to the greatest extent and adopted spread areas roughly 4-fold larger than any of the other cell lines. Interestingly, the spread area did not differ dramatically between aligned collagen fibrils assembled on mica and those transferred to glass except for HFFs and HaCaTs, where areas for both cell lines were larger when the collagen fibrils were transferred to the stiff glass substrate (**Figure 2C**). The degree of directional alignment differed dramatically across the cell lines. HFF and MDA-MB-231 cells robustly oriented on aligned collagen fibrils, whether the collagen fibrils were assembled on mica or transferred to glass (**Figure 2D**). WM-266-4 cells oriented well on aligned collagen fibrils assembled on mica, but poorly on those transferred to glass. HaCaTs adopted an intermediate degree of orientation, whereas THP-1 cells adopted a poor degree of orientation on aligned collagen fibrils. Both HaCaTs and THP-1s did not show differences across the two substrates. The trends in aspect ratio were similar to those in directionality index, indicating that aspect ratio is a reasonable proxy for contact guidance ability (**Supplementary Figure 2**). Consequently, mesenchymal cells (HFFs and MDA-MB-231s) were best at orienting on aligned collagen fibrils, followed by cells that adopt an epithelial morphology (HaCaTs and WM-266-4s). Amoeboid cells (THP-1s) were worst at orienting on aligned collagen fibrils. Furthermore, there are substrate-specific differences in responses to aligned collagen. HFFs and HaCaTs were better spread on the stiff substrate (glass), whereas MDA-MB-231 and WM-266-4 cells were better aligned on the soft substrate (mica).

**Figure 2.**
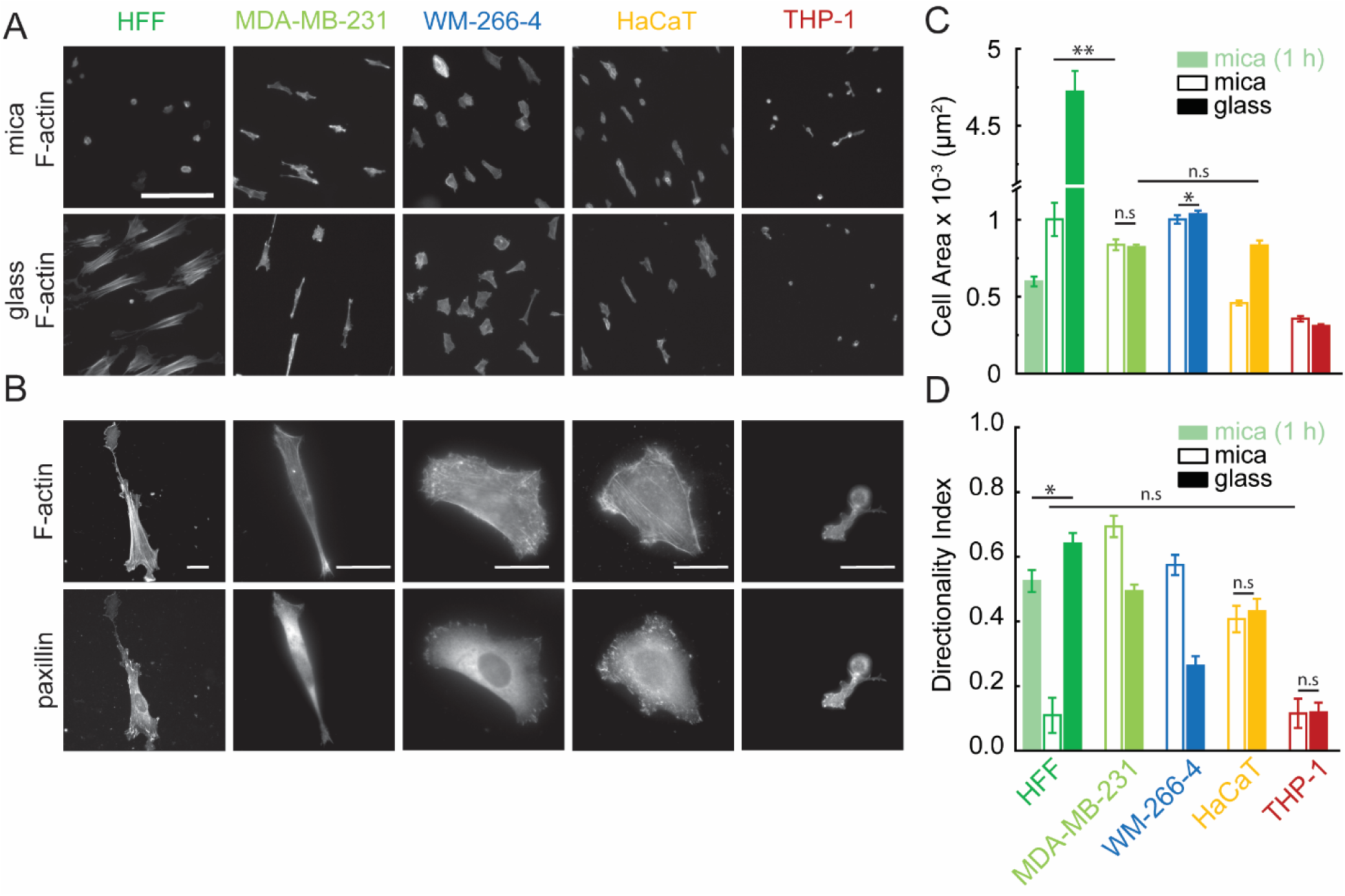
Different cells spread and align to different extents on aligned collagen fibrils. (**A**) Images of F-actin staining of cells on both mica (top) and glass (bottom). Scale bar is 200 μm. (**B**) High resolution image of F-actin and paxillin staining of cells on glass. Scale bars are 25 μm. Cell (**C**) area (*N_samples_* ≥ 4 and *N_cells_* ≥ 304) and (**D**) Directionality index (*N_samples_* ≥ 4 and *N_cells_* ≥ 304) of different cell lines plated on collagen fibrils on mica (open bars) or those transferred to glass (solid bars). Statistical significance was assessed by comparing mica and glass conditions as well as each cell line. All comparisons showed statistically significant differences with *p* ≤ 0.001, except for the comparisons explicitly marked on the plot. Error bars represent 95% confidence intervals.

Given that some cell lines directionally migrate better on collagen fibrils chemically attached to glass substrates, while others prefer collagen fibrils that are loosely attached to mica, we explored the role of both external mechanical properties of the substrate and internal contractile force generation properties of the cell lines. Many of the cell lines showed measurable, but modest dependence of directionality on stiffness, except for WM-266-4 cells, which showed a large and steadily declining directionality for increasing stiffness (**Figure 3A**). We observed a different stiffness for maximal directional alignment across the cell lines. HFFs appeared to align best on 0.2 kPa, MDA-MB-231 and WM-266-4 cells aligned best on mica and HaCaTs aligned best on 2 kPa. The poor directionality of THP-1 cells hindered determining the stiffness of optimal directional migration. Areas and aspect ratios were roughly the same for the cells across stiffnesses, except for HFFs and HaCaTs, which showed robust increases in area as stiffness increased (**Supplementary Figure 3A-C**). What accounts for these contact guidance differences among cells? To test whether differences in receptor expression explains this, we measured the expression level of integrins and discoidin domain receptors (DDR), two families of collagen receptors (**Supplementary Figure 4**). There was a modest, but statistically insignificant Pearson’s correlation coefficient between directionality index on glass and β1-, α1-, α11- and total α-integrin expression. All other collagen receptors showed poor correlation between directionality index and receptor expression.

**Figure 3.**
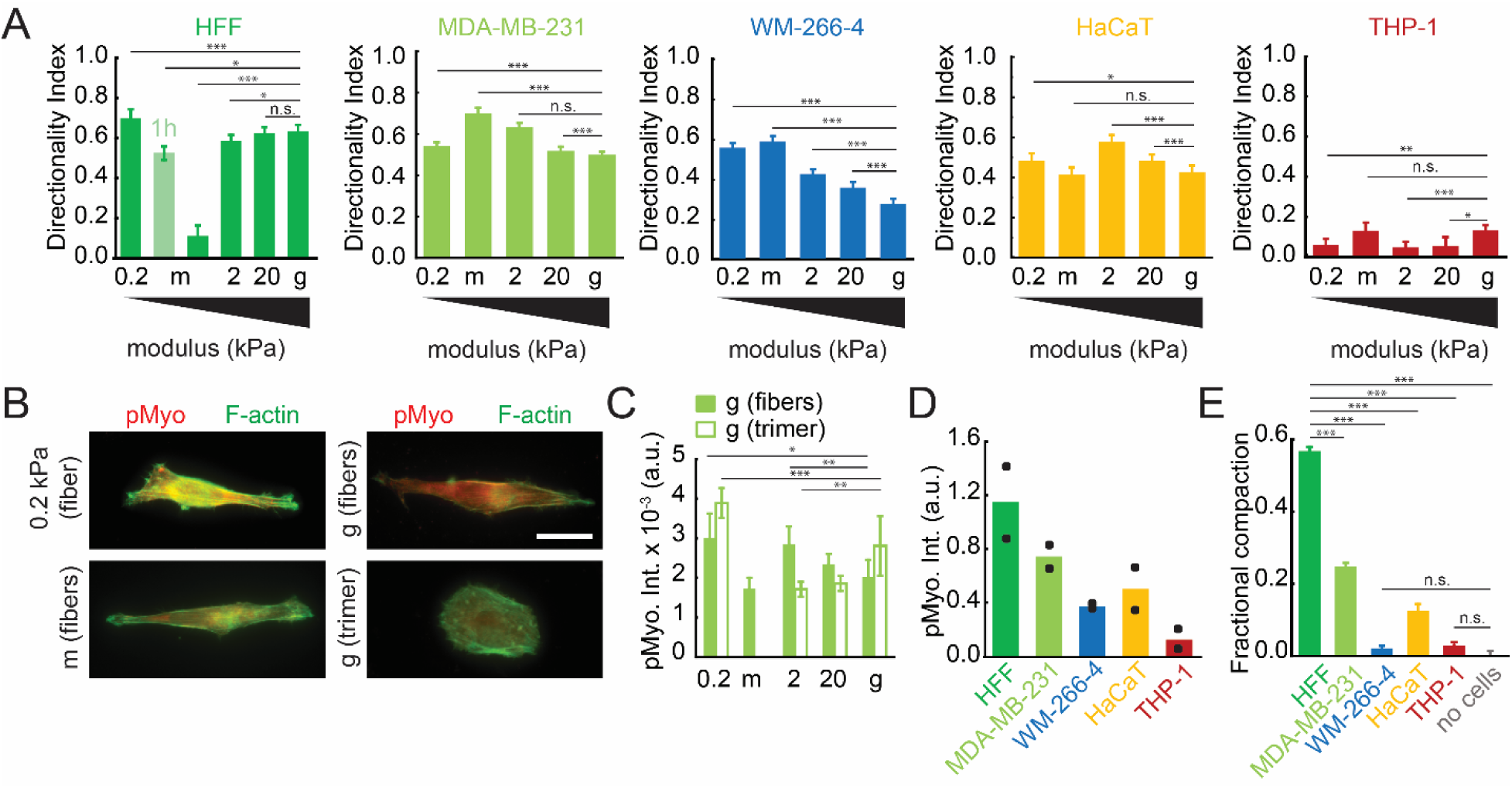
Higher contractile cells engage in optimal contact guidance at lower stiffness. (**A**) Directionality index of various cell lines on different substrates, including 0.2 kPa PAA (0.2), mica (m), 2 kPa PAA (2), 20 kPa PAA (20) and glass (g). *N_samples_* ≥ 4 and *N_cells_* ≥ 304. Statistical significance was assessed by comparing all conditions to the glass condition, where * denotes *p* ≤ 0.05, ** denotes *p* **≤** 0.01 and *** denotes *p* ≤ 0.001. (**B**) F-actin (green) and phosphomyosin (pMyo, red) localization in MDA-MB-231 cells on 0.2 kPa PAA gel with transferred fibrils, glass (g) with transferred fibrils, mica (m) with aligned fibrils, and glass (g) with deposited collagen trimers. The scale bar is 25 μm. (**C**) Phosphorylated myosin (pMyo) intensity in immunofluorescence images of fixed MDA-MB-231 cells on various substrates, including 0.2 kPa PAA gel (0.2), mica (m), 2 kPa PAA (2), 20 kPa PAA (20), and glass (g) with both collagen fibrils (solid bars) and trimers (open bars), *N_samples_* ≥ 2 and *N_cells_* ≥ 28. Statistical significance was assessed by comparing all conditions to the glass condition. (**D**) Phosphorylated myosin (pMyo) intensity from western blots. *N_samples_* ≥ 2. (**E**) Fractional compaction calculated from initial and 48 hours diameters of collagen gels across all cell lines. *N_samples_* ≥ 3. Statistical significance was assessed by comparing all conditions to the HFF condition. Statistical significance was also assessed by comparing all conditions to the no cell condition. For this only the insignificant (n.s.) comparisons are shown. Error bars represent 95% confidence intervals.

Likely, differences in cytoskeletal control and not receptor expression explain differences in contact guidance. Indeed, studies have indicated that myosin activity regulates contact guidance.^[8,13,21,39,40]^ We hypothesized that cells with higher directionality would exhibit higher levels of phosphorylated myosin (p-Myo). Interestingly, cells with roughly similar shapes showed variability in phosphorylated myosin level (**Figure 3B**). Also, conditions that produced the highest directionality had the lowest phosphorylation level (mica) (**Figure 3C**). This differed somewhat from cells on a non-directional collagen cue (collagen trimers on glass), where there appeared to be an increase in phosphorylated myosin as has been seen previously. Cells plated on collagen trimers attached to 0.2 kPa PAA (Polyacrylamide) gel were essentially unspread and showed extremely high signal. However, it is important to note that high phosphorylated myosin level is only efficacious if coincident with a contractile F-actin structure and adhesion to the substrate. Challenges in accurate quantification of immunofluorescence images led to our assessing myosin phosphorylation and contraction by two different means: western blotting and collagen gel compaction. There was a strong and statistically significant Pearson’s correlation coefficient between directionality index on glass and myosin phosphorylation (**Figure 3D and Supplementary Figure 3D**).

To further assess the correlation between contact guidance and contractility, we conducted a compaction test. Cells compact collagen gels at different rates (**Supplementary Figure 3E**). The fractional compaction can be defined as the ratio of gel diameter change relative to the initial diameter over 48 hours. There was a similarly high and statistically significant Pearson’s correlation coefficient between directionality index on glass and compaction (**Figure 3E**). Consequently, control over the F-actin cytoskeleton and not receptor expression appears to explain differences in contact guidance among cells.

### 2.3 Formins and the Arp2/3 complex exert reciprocal control over contact guidance across two distinct cell lines

In order for myosin to contract the F-actin cytoskeleton, F-actin must be polymerized. Two important regulators of F-actin polymerization are formins and the Arp2/3 complex. Formins polymerize linear F-actin, whereas the Arp2/3 complex polymerizes branched F-actin. Aligned stress fibers composed of linear F-actin likely positively regulate contact guidance,^[21]^ whereas broad lamellipodia formed by branched F-actin networks likely give the cell the ability to make turns during migration. Both linear F-actin and branched F-actin draw from the same G-actin pool, creating a tempting hypothesis that when one type of network is diminished, the other type of network is enhanced.^[34]^ Consequently, we hypothesized that these F-actin regulators reciprocally control contact guidance. To test this hypothesis, we employed pharmacological agents including SMIFH2, a formin inhibitor, and CK-666, an Arp2/3 inhibitor.

Inhibition of formins with SMIFH2 in HFF and MDA-MB-231 cells, resulted in dramatic morphology changes (**Figure 4A**). Cells were smaller (**Supplementary Figure 5A&C**) and had diminished stress fiber formation and smooth edges suggestive of branched F-actin networks. Similar morphological responses were seen on both aligned collagen fibrils on mica and aligned collagen fibrils transferred to glass. In addition, cells treated with SMIFH2 showed marked decrease in directionality index (**Figure 4B&C**) and aspect ratio (**Supplementary Figure 5B&D**) at high concentrations. The decrease in directionality of cells plated on aligned collagen fibrils transferred to glass is small, but measurable at 20 μM, increasing through 50 μM, whereas cells plated on aligned collagen fibrils assembled on mica seems to require higher SMIFH2 concentrations to achieve measurable changes in directionality. Given the correlation observed between directionality and contractility and that SMIFH2 is known to block myosin activity at high concentrations (> 50 μM),^[41]^ we measured contractility in two different ways. First, we plated cells on collagen fibrils assembled on mica with and without 20 μM SMIFH2 and measured the kinetics of fibril delamination (**Figure 4D**). There was no statistically significant difference in delamination of collagen fibrils. This suggests that contractility at this concentration was not dramatically inhibited. Compaction kinetics showed a different response (**Supplementary Figure 5E**). Both 20 and 50 μM diminished compaction as compared to untreated cells (**Figure 4E**). Consequently, inhibiting formins diminishes stress fiber formation and directionality at concentrations that likely do not affect myosin activity. At higher formins inhibitor concentrations, myosin is likely inhibited, resulting in dramatic alterations in directionality as we have demonstrated previously.^[42]^ Interestingly, formin inhibition more robustly affects compaction, even at low concentrations. Taken together, this data indicates that formin-mediated F-actin cytoskeletal structures enhance contact guidance.

**Figure 4.**
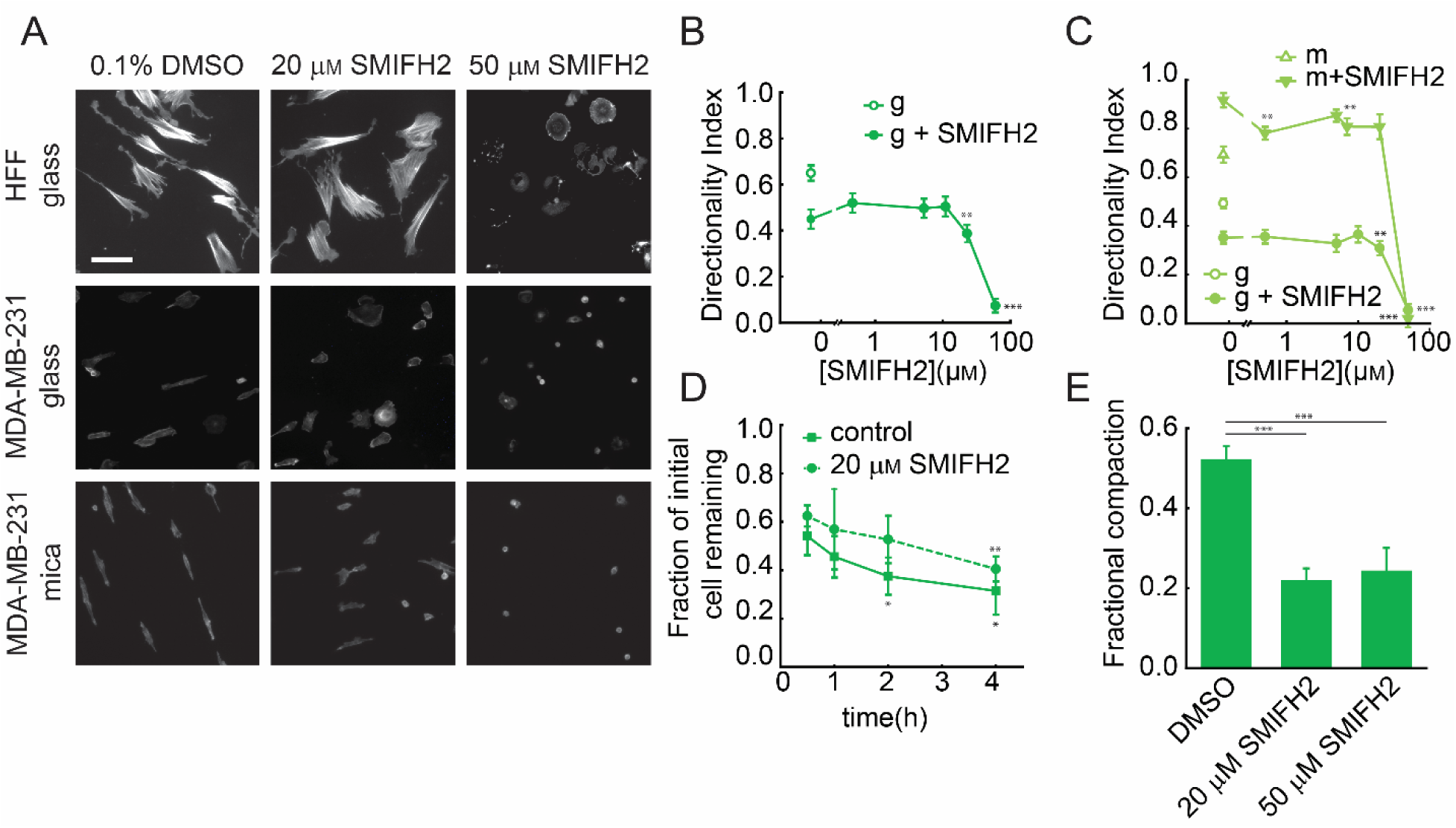
Formin inhibition marginally disrupts mesenchymal cell alignment and collagen fibril delamination, but dramatically diminishes gel compaction. (**A**) Immunofluorescence images of F-actin localization in HFF and MDA-MB-231 cells treated with 0.1% DMSO, 20 µM SMIFH2 and 50 µM SMIFH2 on both mica (m) and glass (g) substrates. The scale bar is 100 µm. (**B**) Directionality index of HFF cells treated with varying concentrations of SMIFH2 on glass (g) substrates, *N_samples_* ≥ 3 and *N_cells_* ≥ 684. The open circles at 0 μM SMIFH2 are cells without any treatment and the solid circles are cells treated with 0.1% DMSO. Statistical significance was assessed by comparing 0.1% DMSO (control) and other treatment conditions. Comparisons not shown were not statistically significant. (**C**) Directionality index of MDA-MB-231 cells treated with different concentrations of SMIFH2 on both mica (m) and glass (g) substrates, *N_samples_* ≥ 2 and *N_cells_* ≥ 168. Comparisons shown are between 0.1% DMSO (control) and other treatment conditions. Comparisons not shown were not statistically significant. (**D**) Fraction of initial HFF cells treated with and without 20 µM SMIFH2 remaining on the mica surface over time as assessed after fixation, *N_samples_* ≥ 4, *N_cells_* ≥ 304 and *N_images_* ≥20. Statistical significance was assessed by comparing to the 0.5-hour time point using a one-way ANOVA (single factor). Significant differences are indicated. No statistical significance was observed between control and 20 µM SMIFH2 for any timepoint. (**E**) Fractional compaction of HFF cells treated with different concentrations of SMIFH2 calculated from initial and 48 hour diameters of collagen gels, *N_samples_* ≥ 3. Statistical significance was assessed between all pairwise comparisons. Comparisons not shown were not statistically significant. Error bars represent 95% confidence intervals.

If our hypothesis is true and there is reciprocal regulation between formins and Arp2/3 over contact guidance, we would predict that decreases in Arp2/3 activity would increase contact guidance. Inhibition of Arp2/3 with CK-666 in HFFs resulted in subtle morphological changes (**Figure 5A**). However, treatment with CK-666 resulted in increased directionality compared to control cells, indicating that Arp2/3 activation inhibits contact guidance (**Figure 5B**). We assessed whether Arp2/3 inhibition could affect the cells ability to delaminate collagen fibrils. It did not, suggesting myosin activity was not greatly altered (**Figure 5C**). Additionally, inhibiting Arp2/3 showed only a small decrease in gel compaction (**Figure 5D** and **Supplementary Figure 6**). Even if Arp2/3 inhibition were to affect compaction through diminishing myosin contractility, the result would be to diminish contact guidance. However, this was not the case as there was a measurable increase in directionality under Arp2/3 inhibition. Consequently, Arp2/3 acts to inhibit contact guidance through the production of branched F-actin networks that give cells an ability to change directions by partially ignoring contact guidance cues. Taken together with data above, formins enhances contact guidance whereas Arp2/3 diminishes contact guidance resulting in reciprocal regulation in HFFs.

**Figure 5.**
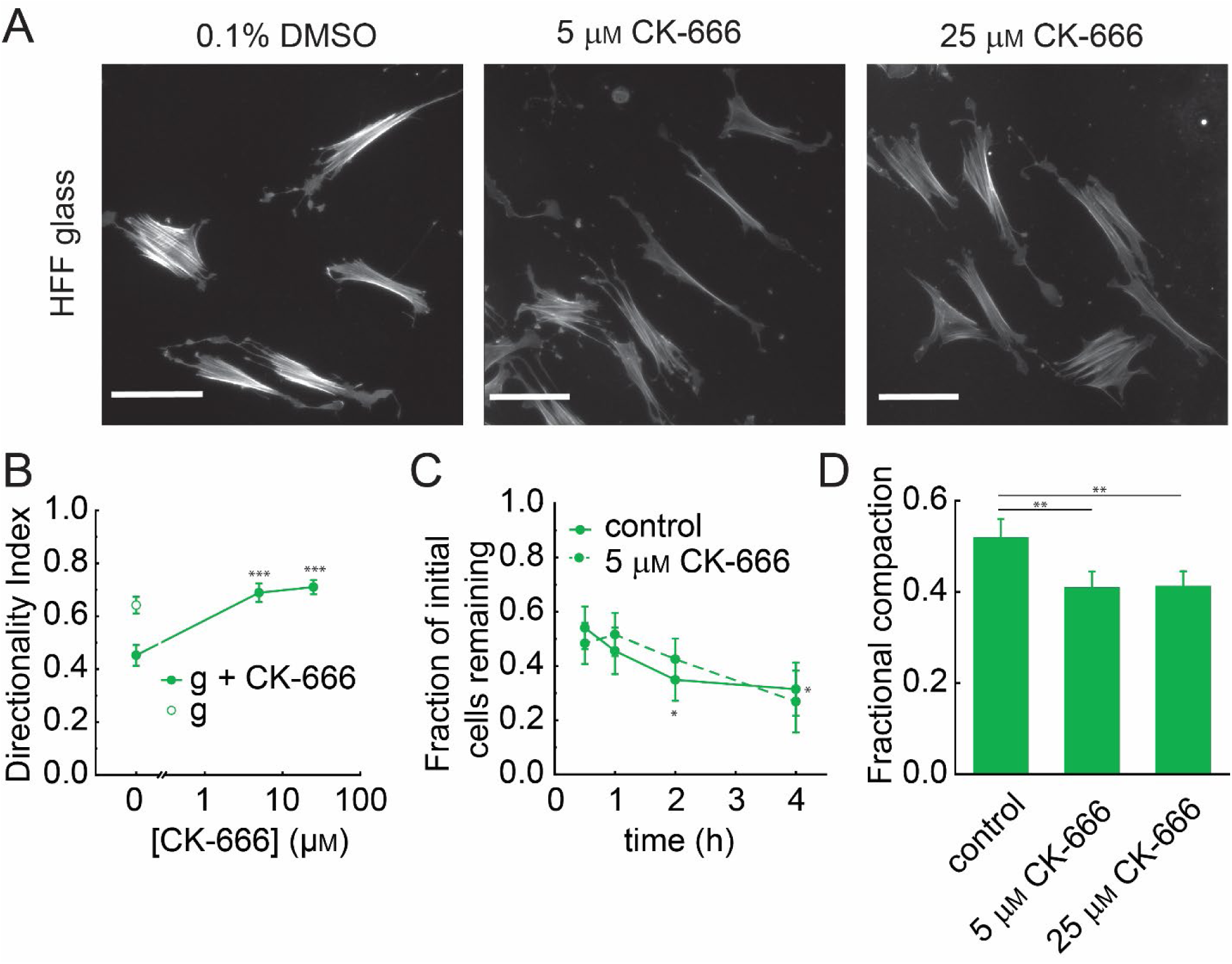
Arp2/3 inhibition enhances mesenchymal cell alignment, but does not robustly affect collagen fibril delamination or collagen gel compaction. (**A**) Immunofluorescence images of F-actin localization in HFF cells treated with 5 µ_M_ and 25 µ_M_ of CK-666 fixed on glass (g). The scale bar is 100 µm. (**B**) Directionality index of fixed HFF cells on glass substrates as a function of CK-666 concentration including 5 and 25 µ_M_ CK-666. The open circles at 0 μM CK-666 are cells without any treatment and the solid circles are cells treated with 0.1% DMSO, *N_samples_* ≥ 3 and *N_cells_* ≥ 594 . Statistical significance was assessed by comparing 0.1% DMSO (control) and other treatment conditions. Comparisons not shown were not statistically significant. (**C**) Fraction of initial HFF cells treated with or without 5 µ_M_ CK-666 remaining on the mica surface over time as assessed after fixation, *N_samples_* ≥ 3, *N_cells_* ≥ 264 and *N_images_* ≥ 20. Statistical significance was assessed by comparing to the 0.5-hour time point using a one-way ANOVA (single factor). Significant differences are indicated. No statistical significance was observed between control and 5 µM CK-666 for any timepoint. (**D**) Fractional compaction of HFF cells treated with different concentrations of CK-666 calculated from initial and 48 h diameters of collagen gels across all cell lines, *N_samples_* ≥ 3. Statistical significance was assessed between all pairwise comparisons. Comparisons not shown were not statistically significant. Error bars represent 95% confidence intervals.

To test whether this reciprocal control of contact guidance by formins and Arp2/3 exists in other cell lines, particularly those with distinct migration behaviors, we examined contact guidance in HaCaTs (**Figure 6A**). Both SMIFH2 and CK-666 modestly affected cell area, but large affects were not observed until relatively high concentrations, where SMIFH2 decreased area and aspect ratio and CK-666 increased area and aspect ratio (**Figure 6B**). Directionality index showed a different response. At low concentrations of SMIFH2, the directionality index of HaCaTs was dramatically diminished as was seen in HFFs (**Figure 6D**). Interestingly, directionality index increased at higher concentrations of SMIFH2 (10 μM) and matched the response in aspect ratio (**Figure 6C**). As SMIFH2 concentration is increased to high levels, myosin contractility is likely inhibited, resulting in a decrease in contact guidance and diminished directionality index and aspect ratio. CK-666 had the opposite effect. At low concentrations of CK-666 (2 μM), the directionality index of HaCaTs is enhanced as was seen in HFFs. However, as the concentration of CK-666 is increased, contact guidance diminished to control levels. This suggests that Arp2/3 activity diminishes contact guidance, but some Arp2/3 activity is needed for high contact guidance in HaCaTs. Furthermore, when both inhibitors (SMIFH2 and CK-666) were added simultaneously, intermediate levels of directionality index were observed. Overall, the contact guidance response to formin inhibition seems to mirror the response to Arp2/3 inhibition. Taken together formins and Arp2/3 seems to exert reciprocal control over contact guidance across two cells with very different migration behaviors.

**Figure 6.**
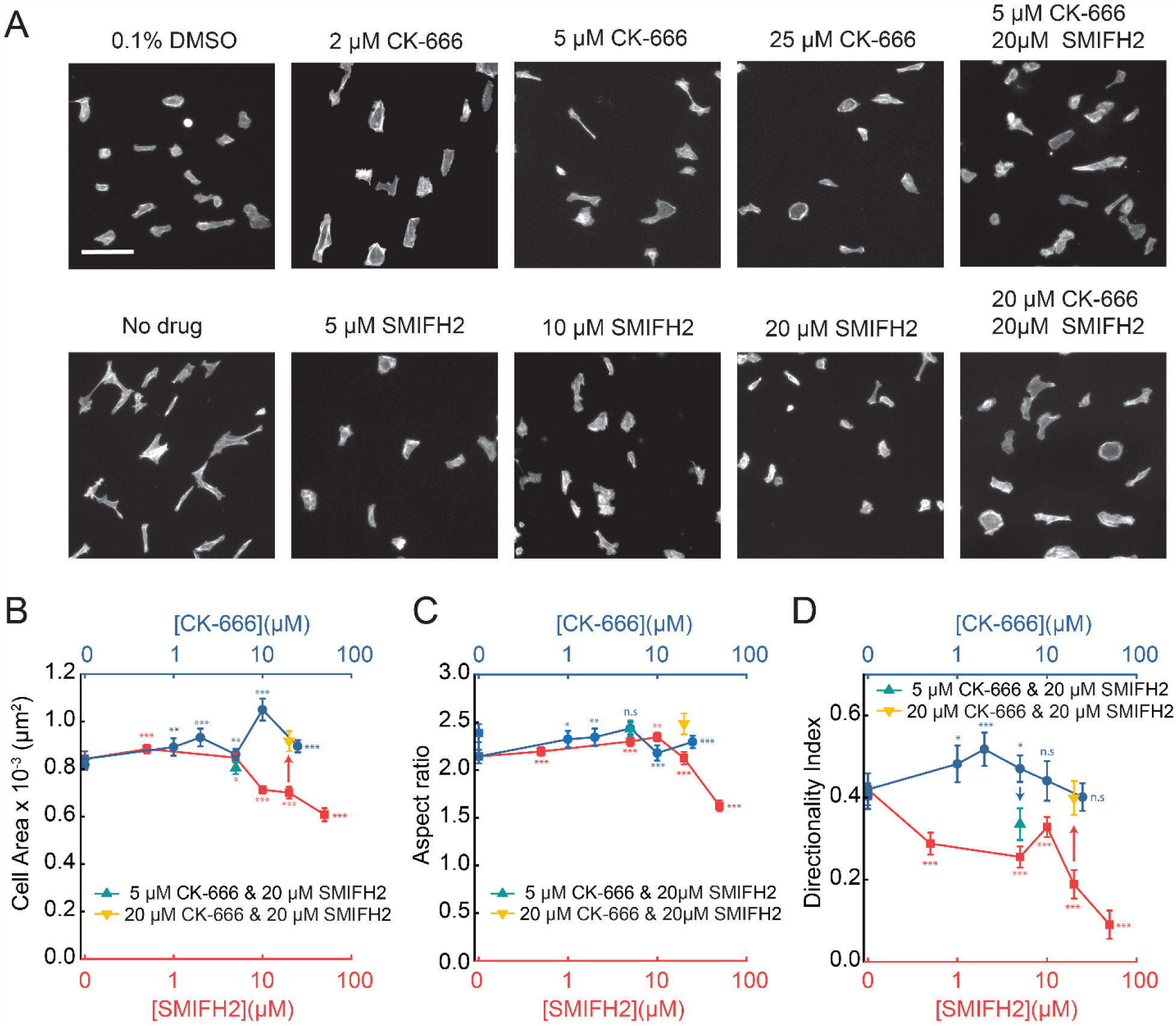
Formin and Arp2/3 inhibition result in reciprocal changes in directionality. (**A**) Immunofluorescence images of F-actin localization in HaCaT cells treated under various conditions: no drug, 0.1% DMSO, CK-666 (2 µM, 5 µM and 25 µM), SMIFH2 (5 µM, 10 µM and 20 µM), and combinations of CK-666 and SMIFH2 (5 µM + 20 µM; 20 µM + 20 µM). The scale bar is 100 µm. (**B**) Cell area, **(C)** aspect ratio and **(D)** directionality index across treatment conditions. *N_samples_* ≥ 2 and *N_cells_* ≥ 504. Statistical significance was assessed by comparing 0.1% DMSO (control) and other treatment conditions. Error bars represent 95% confidence intervals.

### 2.4 The switch-like directional migration behavior of HaCaTs is reciprocally modulated by formins and Arp2/3

We established that formins and the Arp2/3 complex reciprocally regulate contact guidance in both HFFs and HaCaTs. HFFs robustly engage in contact guidance on aligned collagen fibrils. HaCaTs on the other hand seem to be less robust in their alignment (**Figure 2 & 3**). However, it is difficult to determine the direction of HaCaT cell migration in fixed images, due to the fact that they are less aligned than HFFs.

Consequently, in order further explore the role of reciprocal regulation of contact guidance by formins and the Arp2/3 complex we examined live cell migration of HaCaTs. We examined HaCaT migration on aligned collagen fibrils assembled on mica or transferred to 2 kPa PAA gel or glass. Montages of live cell migration showing images at 0, 2, 4, and 8 hours on these substrates revealed interesting behavior (**Figure 7A**). Unlike mesenchymal cells which migrate solely parallel to collagen fibril alignment, HaCaT cells migrated both parallel to the collagen fibrils as well as perpendicular to collagen fibrils, exhibiting a switching behavior (**Figure 7A**). Both parallel and perpendicular movement complicates interpretation of contact guidance from fixed cells. Consequently, we assessed directionality index measured either morphologically (fixed) or by using the angle of the velocity vector during migration (live). Directionality index was lower on both collagen fibrils assembled on mica and those transferred to 2 kPa PAA when measured by migration as compared to morphology (**Figure 7B**). Directionality index increased with substrate stiffness, progressing from mica to glass (**Figure 7B**). Interestingly, cell speed exhibited an inverse trend (**Figure 7C**), demonstrating that stiffer substrates slow down cell migration but enhance its directedness. In order to understand the reason for changes in directionality, given migration both parallel and perpendicular to collagen alignment, we measured the residence time that a cell remains in either parallel (migration angle < 30° with respect to the aligned fibril direction) or perpendicular (migration angle > 60° with respect to the aligned fibril direction) movement. While HaCaT cells spend more time migrating parallel to the collagen fibrils, they do spend a considerable amount of time migrating perpendicularly (**Figure 7D**). To our knowledge, this is a yet to be characterized contact guidance phenomenon. HaCaTs spend more time in parallel movement on stiffer substrates (collagen fibrils transferred to glass) than on softer substrates (collagen fibrils assembled on mica). This agrees with the data showing that HaCaTs migrating on collagen fibrils transferred to glass have a higher directionality index. In addition to residence time, we quantified the fraction of the time cells migrate in a parallel or perpendicular manner (**Figure 7E**). Increased stiffness diminished perpendicular movement compared to parallel movement, but migration on the 2 kPa PAA substrate showed the most evenly balanced parallel and perpendicular movement. Consequently, unlike HFFs and MDA-MB-231 cells, HaCaTs have two modes of directed migration on aligned collagen fibrils, one that is parallel and one that is perpendicular. Furthermore, increasing the stiffness of the underlying substrate biases parallel migration whereas softer substrates, including fibers assembled on mica promote more perpendicular movement.

**Figure 7.**
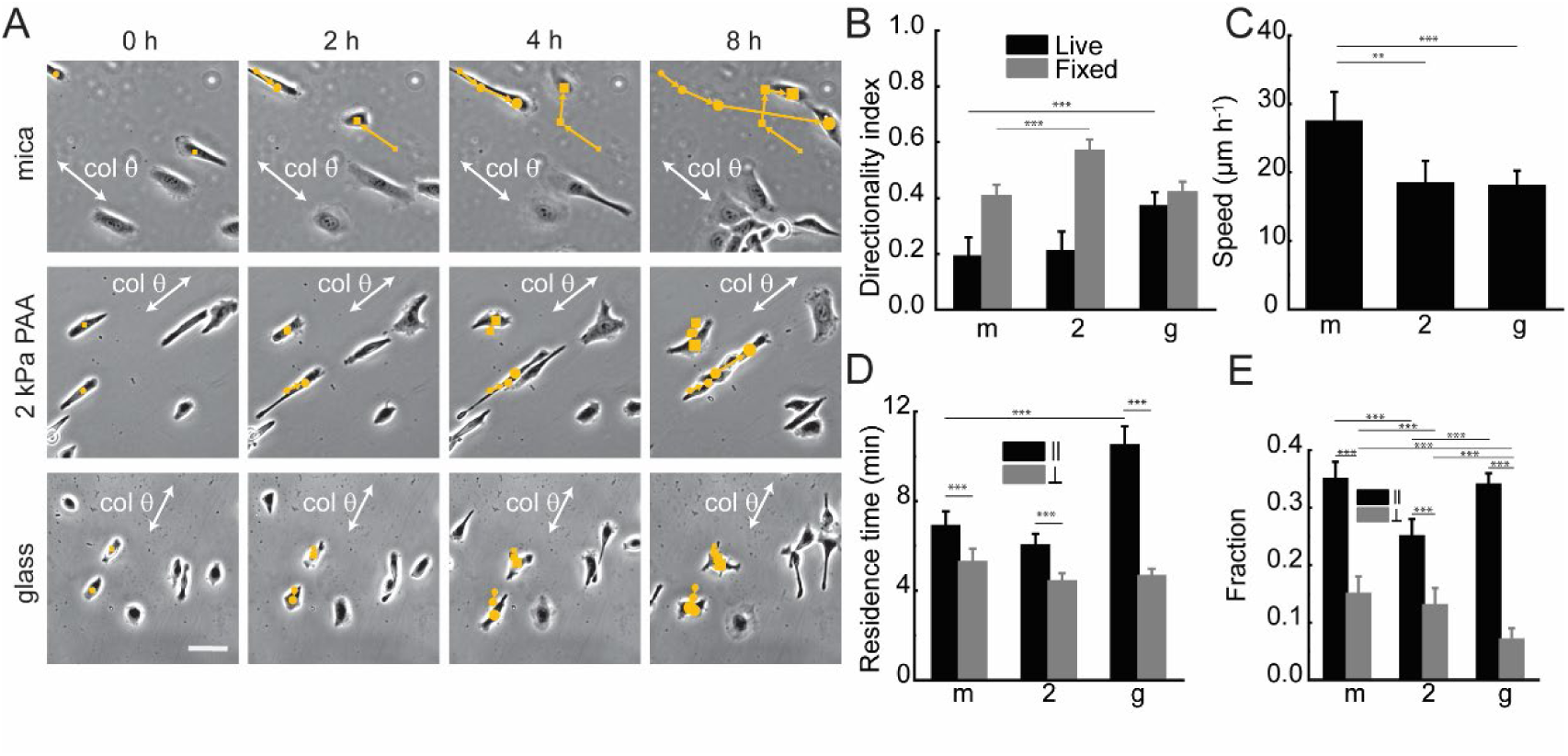
HaCats exhibit both parallel and perpendicular movement that generates low directionality. (**A**) Phase contrast, live cell migration montages of HaCaT cells on different substrates: mica (m), 2 kPa PAA (2), and glass (g). Circles and squares mark previous cell positions. The collagen fibril direction is denoted with the white arrow (col θ). The scale bar is 50 μm. (**B**) Directionality index and (**C**) cell speed, fixed cells: *N_samples_* ≥ 3 and *N_cells_* ≥ 561, and live cells: *N_samples_* ≥ 4 and *N_cells_* ≥ 41. Statistical significance in directionality index was assessed by comparing fixed cells with other fixed cell groups and live cells with other live cell groups for each substrate. Comparisons not shown were not statistically significant. Statistical significance in speed was assessed between all pairwise comparisons. Comparisons not shown were not statistically significant. (**D**) Residence time, indicating the duration cells spend in parallel or perpendicular migration with respect to collagen fibrils and (**E**) fraction of migration steps taken by cells in parallel and perpendicular directions relative to collagen fibrils, *N_samples_* ≥ 4 and *N_cells_* ≥ 41. Statistical significance was assessed by comparing parallel steps between different conditions and comparing perpendicular steps between different conditions as well as comparing parallel and perpendicular steps for each condition. Comparisons not shown were not statistically significant. Error bars represent 95% confidence intervals.

Given that HaCaT cells have intermediate directionality and that they have both parallel and perpendicular movement, we were interested in assessing the role of formins and Arp2/3 in controlling directionality and the balance of these different directional migration modes. As demonstrated in the previous section, formins enhance directionality, whereas Arp2/3 diminishes directionality. A hypothesis that flows from this observation is that formins enhance parallel movement, whereas Arp2/3 enhances perpendicular movement. HaCaTs were imaged during migration. Montages are shown containing images taken at 0, 2, 4, and 8 hours (**Figure 8A**). As described in **Figure 6**, formin inhibition appears to diminish contact guidance and Arp2/3 inhibition seems to enhance contact guidance. We quantified directionality index and speed. When fibers were transferred to glass, directionality index assessed by fixed cell morphology and directionality index assessed using live cell migration tracks are essentially the same (**Figure 8B**). Directionality index decreased robustly when formins were inhibited as we described above. Inhibition of Arp2/3 modestly reduced directionality. Speed showed a different response. Cells under formin or Arp2/3 inhibition moved at the same rate as each other as well as faster than controls (**Figure 8C**). Since HaCaT cells move in both parallel and perpendicular fashions, we assessed whether inhibiting formins or Arp2/3 differentially affected these different directional modes of migration. Similar to our previous analyses (**Figure 7**), we quantified and analyzed cell movements to evaluate the effects of formins or Arp2/3 inhibitors. Decreases in residence time (**Figure 8D**) tracked with changes in directionality index (**Figure 8B**). For cells treated with the Arp2/3 inhibitor (25 µ_M_ CK-666), we observed an increase in both parallel and perpendicular steps (**Figure 8E**), resulting in a slightly lower directionality index, compared to the control due to the balanced distribution between parallel and perpendicular movements (**Figure 8B**). In contrast, cells treated with the formin inhibitor (20 µ_M_ SMIFH2) showed a similar fraction of parallel steps compared to the control but an increased fraction of perpendicular movements, resulting in a markedly lower directionality than the control. These live-cell contact guidance results for HaCaTs agree broadly with the fixed-cell contact guidance results for HaCaTs, HFFs and MDA-MB-231s shown above. Formins tend to enhance contact guidance, whereas Arp2/3 diminishes contact guidance.

**Figure 8.**
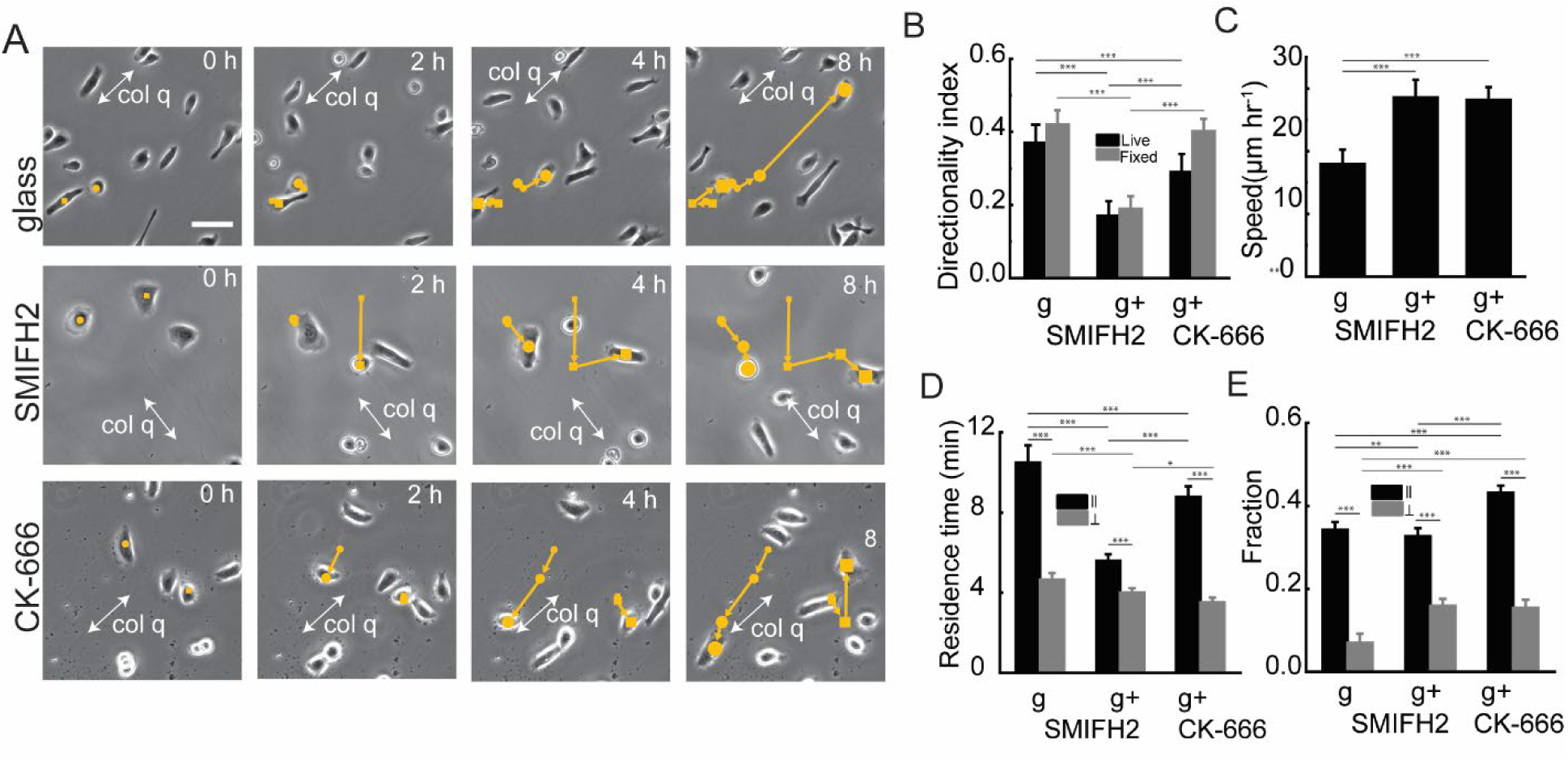
In HaCaTs Arp2/3 inhibition increases parallel and perpendicular migration, whereas formins inhibition increases perpendicular migration only. (**A**) immunofluorescence images of HaCaT cells fixed on glass treated with 0.1% DMSO, 20 µ_M_ SMIFH2 and 25 µ_M_ CK-666. Scale bar is 50 µm (**B**) Live cell migration montages of HaCaT cells on glass without inhibitors, with 20 µ_M_ SMIFH2 and with 25 µ_M_ CK-666. Scale bar is 50 µm (**C**) Directionality index of both fixed *N_samples_* ≥ 4 and *N_cells_* ≥ 716 and live cells .*N_samples_* ≥ 5 and *N_cells_* ≥ 85 under different conditions: no inhibitor, 20 µ_M_ SMIFH2 and 25 µ_M_ CK-666. Statistical significance in directionality index was assessed by comparing fixed cells with other fixed cell groups and live cells with other live cell groups for each substrate. Comparisons not shown were not statistically significant. Statistical significance in speed was assessed between all pairwise comparisons. Comparisons not shown were not statistically significant. (**D**) Cell speed calculated from live cell migration under the same conditions: no inhibitor, 20 µ_M_ SMIFH2 and 25 µ_M_ CK-666, *N_samples_* ≥ 5 and *N_cells_* ≥ 85 (**E**) Average residence time of cells moving parallel (less than 30° deviation) or perpendicular (more than 60° deviation) to collagen fibrils. *N_samples_* ≥ 5 and *N_cells_* ≥ 85 (**F**) Fraction of cells moving parallel (deviation less than 30°) or perpendicular (deviation more than 60°) to collagen fibrils. *N_samples_* ≥ 5 and *N_cells_* ≥ 85. Statistical significance was assessed by comparing parallel steps between different conditions and comparing perpendicular steps between different conditions as well as comparing parallel and perpendicular steps for each condition. Comparisons not shown were not statistically significant. Error bars represent 95% confidence intervals

## 3. Discussion

In this paper we leverage an approach to assemble collagen fibrils on mica in an aligned fashion. We have previously shown that aligned collagen fibrils assembled on mica replicate contact guidance differences across cells in 3D systems ^[13]^ that are masked in systems like grooves or microcontact printed lines.^[21]^ However, highly contractile cells delaminate the collagen fibrils, complicating the assessment of contact guidance. Consequently, we optimized an approach to transfer these collagen fibrils to functionalized substrates, including those with tunable stiffness. This transfer approach allowed us to overcome the challenge of highly contractile cells and characterize contact guidance across cells with different contractile abilities. This is important because, while others have shown that there are cell type differences in response to contact guidance cues,^[27–30]^ these previous reports have been conducted on grooves, which mask cell-type differences in contact guidance.^[21]^ Consequently, our system can leverage advantages seen in 2D systems, including easier imaging, while retaining key predictive abilities including cell-type differences in contact guidance.

The collagen fibril assembly and transfer system not only blocks delamination of collagen fibrils by highly contractile cells, it also allows us to probe extrinsic factors that drive cell-type differences. Highly contractile cells like HFFs engage in contact guidance that is independent of stiffness. Similarly, in weakly contractile cells like THP-1s, contact guidance is also unaffected by the underlying substrate stiffness. Only cells with intermediate contractile behavior, including MDA-MB-231, WM-266-4 and HaCaT cells show noticeable maxima in directionality index. The fact that different cells respond differently to contact guidance cues and that the response is stiffness-dependent matches intuition about the function of different cells. Fibroblasts must remodel the ECM while moving along fibrils during wound healing. Since stiffness changes dramatically during the wound process, but fibroblasts must continue to contact guide and generate traction force, they may need to be less sensitive to stiffness cues. Similarly, for amoeboid cells, which must randomly sample the wound environment, it might be important for them to ignore the directional cue. Indeed, aligned collagen fibrils in the tumor microenvironment do not seem to affect T-cell infiltration.^[25]^ Substrate stiffness is an appreciated modulator of cell function and likely tunes contact guidance fidelity due to overlapping intracellular mechanisms.^[43,44]^ The literature shows conflicting reports examining contact guidance in MDA-MB-231 cells on flexible grooves. One report indicated that increased stiffness diminishes the cell aspect ratio,^[37]^ indicating that that contact guidance decreases under stiffer conditions, however another report showed increased elongation and small increases in cell alignment in MDA-MB-231 cells.^[30]^ We showed decreases in MDA-MB-231 cell contact guidance in 3D environments after collagen crosslinking leading to increases in stiffness,^[13]^ matching what we observed on aligned fibrils in 2D here, suggesting again that the aligned collagen fibril assembled system mimics 3D environments and provides clarity with respect to discrepancies observed when using groove systems.

One important question to be answered is what causes these cell type differences in contact guidance. To explore this, we examined receptor expression and cytoskeletal regulation. We correlated collagen receptor expression of both integrins and DDRs to contact guidance fidelity on collagen fibrils transferred to glass. DDRs poorly predict contact guidance ability. Many of the individual α-integrins also showed a poor ability to predict contact guidance ability. β1-integrin and total α-integrin when correlated to directionality index, showed modest but statistically insignificant Pearson’s correlation coefficients. We and others have shown that altering β1-integrin activation can tune contact guidance,^[21,45]^ consequently, it appears that integrin expression is needed, but the level does not determine contact guidance ability. Consequently, we explored cytoskeletal regulation as being the driving factor to determine contact guidance ability. Multiple studies underscore the role of contractility in cell alignment and contact guidance.^[8,13,17,21,29,30]^ While early contact guidance stages are contractility-independent, later stages require actomyosin traction forces.^[40]^ Studies on microcontact-printed collagen lines further show that MTLn3 cell directionality is diminished after blocking contractility.^[8]^ In a variety of cells, blocking contractility reduced alignment along microgrooves, weakening contact guidance.^[29,30]^ Similarly, in 3D collagen gels, inhibiting contractility with blebbistatin or a myosin phosphorylation blocker lowered directionality to levels observed on non-aligned substrates in MDA-MB-231 cells.^[13,46]^ While myosin is demonstrated to be important, its role as a universal predictor of contact guidance ability across diverse cell lines has not be established. This study shows that intracellular contractility is a superb predictor of contact guidance behavior. Both myosin phosphorylation and 3D collagen gel compaction assays revealed strong and significant Pearson’s correlation coefficients. Contractile ability better predicts contact guidance on aligned collagen fibrils that are stiff (transferred to glass) as compared to those that can be remodeled (collagen fibrils assembled on mica). Myosin phosphorylation is elevated in environments when cells can recruit fibrils,^[47]^ but we assessed myosin phosphorylation on tissue culture plastic due to the infeasibility of using western blotting on our aligned collagen fibril substrates as our collagen substrates contain a small number of cells compared to the number needed for western blotting. The fact that cells experience similar stiffness on tissue culture plastic as they do on collagen fibrils transferred to glass likely explains the better correlation between myosin phosphorylation and contact guidance on collagen fibrils assembled on glass as compared to the correlation between myosin phosphorylation and contact guidance on collagen fibrils assembled on mica. Consequently, we have shown that not only is myosin contractility important to contact guidance, but the level of myosin contractility can robustly predict the efficiency of contact guidance and that this is a much better predictor of contact guidance than collagen receptor expression.

Since myosin contractility was a good predictor of contact guidance, we were interested in other cytoskeletal regulators that could tune contact guidance ability. Consequently, we explored the regulation of contact guidance by formins and Arp2/3. In 1D contact guidance systems, where cells are confined to 1D migration only, formins are important, whereas Arp2/3 is dispensable in maintaining speed.^[35,48,49]^ However, these 1D systems dimensionally restrict migration in a way that does not allow the assessment of how directional fidelity is controlled. In 2D systems of contact guidance that include microcontact printed lines and ridges, formins seem to play a positive role in controlling directional fidelity.^[36,37]^ The role of Arp2/3 is less clear. Work exploring contact guidance on flexible substrates indicates that inhibiting Arp2/3 diminishes contact guidance ^[37]^ whereas other work on stiff substrates indicates that inhibiting Arp2/3 enhances contact guidance.^[36]^ Consequently, whether formins and Arp2/3 reciprocally regulate contact guidance is not known. Two observations suggest reciprocal regulation. First, the F-actin networks that formins and Arp2/3 create are distinct in nature. Formins generate linear F-actin structures like various forms of stress fibers that help the cell establish and maintain traction in a particular direction of migration through stable focal adhesions attached to the ECM. Arp2/3 generates branched F-actin networks that constitute the lamellipodium, a rounded structure that can also facilitate dynamic adhesion and can bias migration in a particular direction but has much more plastic control of directional persistence. Second, it has been elegantly shown in a variety of contexts that G-actin can be limiting, so diminishing of linear F-actin structures through formin inhibition results in more branched F-actin structures and vice versa, setting up feedback for reciprocal control by these two distinct F-actin networks.^[34,50]^ We show reciprocal regulation of contact guidance by formins and the Arp2/3 complex across two cell lines with drastically different migration strategies (**Figure 9A**). In HFFs, formin inhibition decreases directionality index indicating a diminished contact guidance fidelity, whereas Arp2/3 inhibition increases directionality index indicating an enhanced contact guidance fidelity. This response can be predicted by a simple mathematical model fitted to the experimental data (**Figures 5&6**), whereby branched F-actin and linear F-actin draw from the same pool of G-actin and directionality index is proportional to the concentration of linear F-actin in the cell (**Figure 9A**). When SMIFH2 inhibits assembly of linear F-actin (*IC*_50_ ∼ 15 μM),^[52]^ directionality index decreases. When the concentration of SMIFH2 approaches the *IC*_50_ for myosin inhibition (30 μM),^[41]^ additional decreases in directionality index are seen, due to inhibiting myosin contractility. Conversely, when CK-666 inhibits the Arp2/3 complex, directionality index increases. We speculate that this is due in part to the additional G-actin that is available for use by formins to generate additional linear F-actin, promoting contact guidance. There is a more complex behavior in HaCaTs. Small concentrations of SMIFH2 result in decreases in directionality index, as seen in HFFs. However, larger doses of SMIFH2 (10 μM) increase directionality index. This can be modeled by adding an additional model component, SMIFH2 blocking branched F-actin with cooperativity and a higher *IC*_50_. We speculate that this is accomplished indirectly, through diminished levels of linear F-actin. Indeed, others have shown that formins are important in enhancing lamellipodial strength ^[51]^ and thus cooperate with Arp2/3.^[50]^ Inhibition of that cooperation may diminish branched F-actin concentration, freeing up some G-actin for additional linear F-actin assembly. However, at larger concentrations of SMIFH2, linear F-actin assembly and possibly myosin inhibition diminishes, reducing directionality index. While this is speculative, something additional must be added to the model to explain the reproducible increase in directionality index seen in HaCaTs at intermediate SMIFH2 concentrations (10 μM). Conversely, at low concentrations of CK-666, directionality index increases as seen in HFFs, albeit marginally. However, larger doses of CK-666 (10 μM) diminish contact guidance.

**Figure 9.**
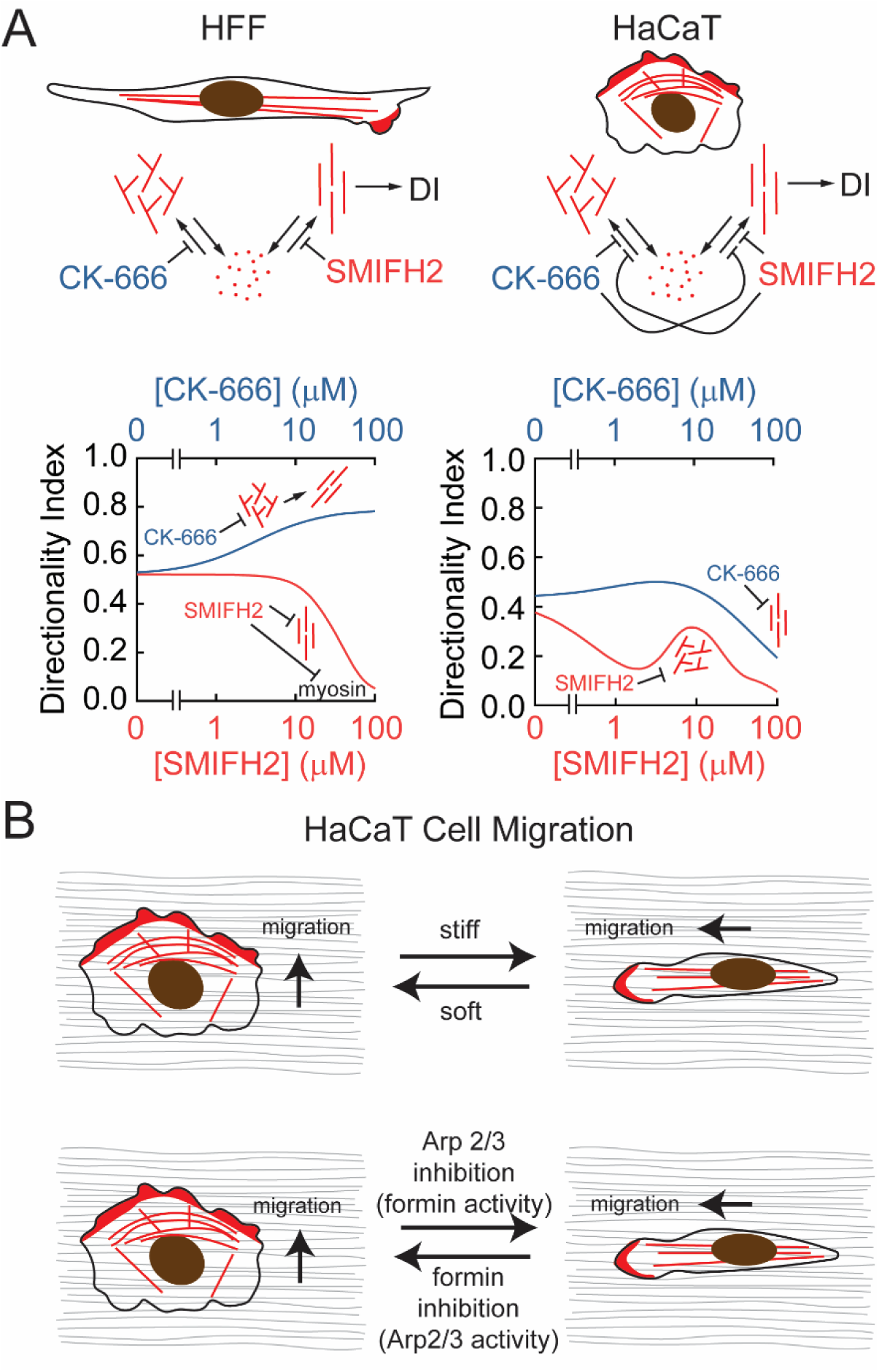
Model of the role of Arp2/3 mediated F-actin branching and formin mediated linear F- actin in directing cells on aligned collagen. **(A)** A simple mass action kinetics model of a shared G-actin pool explains both the decrease in contact guidance when formins are inhibited (SMIFH2) and increase in contact guidance when Arp2/3 is inhibited (CK-666) in HFF cells. At extremely high concentrations of SMIFH2 (> 50 μM), myosin is inhibited, leading to diminished contact guidance. This simple mass action kinetics model does not explain the biphasic behavior in HaCaT cells, where at high CK-666 concentrations, contact guidance decreases and at high SMIFH2 concentrations, contact guidance increases. Inhibition of linear F-actin at high CK-666 concentrations and inhibition of branched F-actin at intermediate SMIFH2 concentrations can explain this behavior. **(B)** HaCaT cells switch between parallel and perpendicular migration behavior on aligned collagen fibers in response to both external cues as well as the balance between Arp2/3 mediated F-actin branching and formin mediated linear F-actin bundles.

Again, this can be modeled by adding an additional interaction, CK-666 blocking linear F-actin with a different *IC*_50_. We speculate that this is accomplished indirectly, through diminished levels of branched F-actin. Branched F-actin could be the source for bundling activity that is observed by some members of the formin family.^[52–54]^ Bundled, aligned F-actin could then be indirectly inhibited through the inhibition of Arp2/3. While the mechanism of this different form of reciprocal regulation of contact guidance in HaCaTs is not currently known, reciprocal regulation across different cell lines does seem to be a reliable feature of formin and Arp2/3 control over contact guidance.

Finally, we have discovered a new mode of migration on aligned collagen fibrils. On strong contact guidance cues like ridges and lines of ECM, all cells respond by migrating parallel to the groove or line direction. As mentioned above our collagen fibril substrates reveal cell type differences that can be masked in these other systems and these differences map to responses observed in 3D. This was demonstrated by the difference between migration of mesenchymal cells (HFFs and MDA-MB-231s), which migrate on collagen fibrils with high fidelity and amoeboid cells (THP-1s), which essentially ignore the collagen fibril direction. We find HaCaTs (epidermal kerotinocytes) can migrate in both a parallel and perpendicular fashion. This behavior is only revealed when examining cell migration and is less apparent when examining cell alignment in fixed images. This appears to be a new phenotype not observed by those examining fixed cell morphology and one that is more readily observable using our aligned collagen fibril system. Normally, during wound healing, collective migration of epidermal keratinocytes occurs resulting in re-epithelialization of the wound. However, we imagine that in some instances single cells detach or are left during injury. As the wound develops, they interact with aligned fibrin or collagen fiber networks that are generated by contractile dermal fibroblasts. It is crucial for these cells to be somewhat agnostic to the directional alignment of the collagen, necessitating both parallel and perpendicular migration. Alternatively, this single-cell multi-directional phenotype might indicate similar collective cell responses on aligned collagen fibrils that has not yet been observed. Indeed, work has been carried out trying to understand collective cell migration on contact guidance cues and sheets indeed prefer migration parallel to the groove, but again, perhaps there are some responses that are masked by these non-fibril systems.^[55–57]^ External signals, such as stiffness, regulate the balance of perpendicular and parallel migration (**Figure 9B**) and both formins and Arp2/3 show reciprocal regulation of contact guidance as assessed through cell migration. This role of formins in diminishing contact guidance and Arp2/3 in enhancing it is similar to what we observed above when examining contact guidance using measures of cell alignment. How the F-actin structure dynamically changes during parallel and perpendicular migration as well as how formins and Arp2/3 regulate this behavior differently during each mode of migration is still an open question and will need to be assessed in the future. The discovery of this additional mode of directional migration will guide the design of the ECM that regulate wound healing and could be encoded into tissue engineered constructs to facilitate directed migration.

## 4. Conclusions

We show that while collagen fibrils assembled on mica produce a useful substrate on which to assess contact guidance. However, highly contractile cells like dermal fibroblasts delaminate collagen fibrils limiting the usefulness of the system. This challenge can be overcome by transferring collagen fibrils to functionalized surfaces, blocking the delamination. We show that contact guidance is dramatically different across cells with different migration modes. Furthermore, these contact guidance differences are not predicted by collagen receptor expression, but rather by control over the cytoskeleton. Myosin phosphorylation and contractile strength correlate well with contact guidance. In addition to myosin contractility, formins and Arp2/3 reciprocally control contact guidance, where formins facilitate better contact guidance and Arp2/3 facilitates poorer contact guidance. Finally, epithelial cells appear to switch between parallel and perpendicular migration on aligned fibrils. This behavior is stiffness-dependent, where stiffer environments result in better contact guidance at slower rates. Formins enhance the parallel movement, whereas Arp2/3 has a richer control over contact guidance, diminishing parallel migration, but also decreasing perpendicular migration. Taken together, this work demonstrates critical aspects of cytoskeletal regulation that prime different cells for efficient contact guidance.

## 5. Materials and Methods

### 5.1 Assembling collagen on mica

A freshly cleaved 15 mm x 15 mm piece of muscovite mica (highest grade VI, Ted Pella, Redding, CA) was used for the epitaxial growth of collagen fibrils. Rat tail collagen type I (Corning, Corning, New York) was diluted using 50 m_M_ Tris-HCl (Fisher Scientific, Waltham, MA) and 200 m_M_ KCl (Fisher Scientific, Waltham, MA) to achieve final concentrations of 10 or 100 μg ml^-1^ and added to the top of the mica. Following the 12-hour incubation period, the mica was washed with nano-pure water. The mica was then placed inside a tissue culture dish, allowed to dry overnight and used the following day. To prepare trimeric collagen structures on mica, 50 m_M_ citric acid-sodium citrate (Fisher Scientific) and 200 m_M_ KCl at pH 4.3 were used to dilute the collagen, and the mica was incubated as above.

### 5.2 Functionalizing and transferring collagen fibrils to glass

First, 22 mm × 22 mm coverslips (Corning, Corning, New York) were treated with piranha solution (3:1 H_2_SO_4_ (Fisher chemical) to H_2_O_2_ (Fisher chemical)) for 1 hour to ensure thorough cleaning and activation. After treatment, the coverslips were immersed in 1% (v/v) 3-aminopropyltriethoxysilane (APTES, Thermo scientific) solution prepared in 1 m_M_ acetic acid (Alfa Aesarfor, Ward Hill, MA) for 2 hours to facilitate silanization. The coverslips were then thoroughly rinsed with deionized water and allowed to air dry. Subsequently, the coverslips were baked in an oven at 100 °C for 1 hour to promote covalent bonding of the APTES layer. Once cooled, the coverslips were treated with 6% (v/v) glutaraldehyde (Electron Microscope Science, Hatfield, Pennsylvania) in Dulbecco’s Phosphate-Buffered Saline (DPBS) for 2 hours to introduce aldehyde functional groups. Following this treatment, the coverslips were extensively washed with deionized water to remove residual glutaraldehyde. Coverslips were good for two weeks. For transferring the collagen, 30% gelatin (Sigma Aldrich, St. Louis, Missouri) was dissolved in water and heated up to 40 °C for an hour, and 500 µl of the gelatin solution was placed on the mica with assembled collagen. After a 1-hour incubation, the gelatin was peeled off and placed on chemically functionalized glass. Following an additional hour of incubation of the gelatin on the target substrate, the samples were washed with 5 ml of DPBS lacking magnesium and calcium (Gibco, Waltham, MA) by placing the DPBS inside the culture dish containing the substrate and incubating at 37°C for 2 hours. After this period, the underside of the sample was cleaned with a Kim wipe (Kim wipe Kimberly-Clark, Irving, Texas), and the substrate was prepared for cell seeding.

### 5.3 Fabricating and transferring collagen fibrils to polyacrylamide gels

First, 22 x 22 mm coverslips were thoroughly cleaned and then covered with 0.1 M NaOH for 10 minutes. Following this, the coverslips were treated with a 5% APTES solution for 5 minutes. The coverslips were then washed three times with deionized water, each wash lasting 5 minutes. Next, a 0.5% glutaraldehyde solution was applied to the coverslips for 30 minutes, followed by additional washes with deionized water. To prepare polyacrylamide gels with different elastic moduli, solutions of 3%, 5%, and 8% acrylamide and 0.03%, 0.1%, and 0.26% bis-acrylamide were prepared and 0.05% Ammonium Persulfate (APS, Bio-Rad, Hercules, California) and 0.15% N,N,N’, N’-Tetramethylethylenediamine (TEMED, Fisher Scientific), respectively, to achieve gels with elastic moduli of 200, 2000, and 20000 Pa ^[58]^.The functionalized coverslips were placed face down onto drops of these solutions on microscope slides, allowing polymerization to occur for 20 minutes. The samples were then treated with 2 mM Sulfo-SANPAH (Thermo Scientific) under UV light for 8 minutes to functionalize their surfaces. The fibril transfer technique was then employed as described in the above section.

### 5.4 Rheological Measurements of Polyacrylamide Gels

To confirm the mechanical properties of the prepared polyacrylamide (PA) substrates, rheological measurements were performed using an Ares G2 equipped with a 25 mm parallel plate rheometer with a parallel plate geometry. Gels were polymerized with varying acrylamide:bis-acrylamide ratios to achieve different cross-linking densities. Frequency sweep tests were conducted at 1% strain across a frequency range of [0.1–100 Hz] to obtain storage modulus (*G*′) and loss modulus (*G*″). The elastic modulus was calculated based on the average *G*′ values at 0.1 Hz from at least 3 independent gel replicates for each condition. As anticipated, a decrease in cross-linker concentration corresponded to a decrease in gel stiffness (**Supplementary Figure 1**).

### 5.5 Characterizing collagen fibrils using Atomic Force Microscope (AFM) imaging and immunofluorescence

The topography of the collagen assembly on mica and transferred to glass was imaged using an AFM (Digital Instruments, now Bruker Nano, Santa Barbara, CA) in tapping-mode with TESPA probes with 40 mV drive amplitude and frequency of 320 kHz. Captured images were processed with the plain fit and flattened routines as appropriate. Collagen fibrils were also stained and imaged using immunofluorescence. Collagen fibril samples were first blocked with blocking buffer (0.02 g ml^-1^ BSA prepared in 10% TBS containing 0.01% Tween-20) for 20 minutes and then incubated with 1:100 collagen antibody (polyclonal rabbit, Thermo fisher, Bio-Rad-2150-1908) for 24 hours. After washing three times with 1x TBS for 5 minutes each, samples were incubated with a 1:200 dilution of donkey anti-rabbit 488 (Jackson Immuno Research, #711-035-152) secondary antibody for 1 hour, followed by additional washes.

### 5.6 Culturing cells and treating with inhibitors

An immortalized human foreskin fibroblast cell line (HFF, ATCC, Manassas, VA), a human mammary carcinoma cell line (MDA-MB-231, ATCC), a metastatic human melanoma cell line (WM-266-4, ATCC), an immortalized human keratinocyte cell line (HaCaT, kind gift from Dr. Torsten Wittmann) and an immortalized monocyte cell line (THP-1, ATCC) were used. MDA-MB-231s, WM-266-4s and HaCaTs were cultured in Dulbecco’s modified Eagles medium (DMEM, Sigma Aldrich). Media was supplemented with 10% fetal bovine serum (FBS) (Gibco, Grand Island, NY), 1% penicillin– streptomycin (Gibco), and 1% Glutamax (Gibco) at 37 °C in 5% CO_2_. The same media with 15% FBS (Gibco) was used for culturing HFF cells. THP-1 cells were cultured in RPMI-1640 media with 10% FBS, 1% penicillin–streptomycin, 0.05 m_M_ BME (β-Mercaptoethanol). When live cells were imaged, cells were placed in culture media Dulbecco’s Modified Eagle Medium that lacked phenol red, but contained 15 m_M_ 4-2-hydroxyethyl) piperazine-1-ethanesulfonic acid (HEPES), 1x Glutamax, 1x penicillin-streptomycin, 10% Fetal bovine serum (FBS). For inhibitor treatments, cells were centrifuged, counted and suspended in culture media. SMIFH2 (Millipore), CK-666 (Millipore) and blebbistatin (Millipore) were dissolved at 50 m_M_, 50 m_M_ and 20 m_M_ stock solutions in dimethyl sulfoxide (DMSO). Stock solutions of the inhibitor were diluted in either culture media or imaging media by at least 1000-fold to make working solutions of 0.5, 5, 10, 20, and 50 m_M_. For the control group, 0.1% DMSO was added to cell culture media or imaging media.

### 5.7 Immunofluorescence imaging and analyses

Cells were plated onto substrates at a concentration of 2 x 10^4^ cells ml^-1^. Cells were fixed and stained to visualize their F-actin structure and quantify their response to contact guidance through morphology and alignment behavior. First, the cells were treated with 4% paraformaldehyde (Fisher Scientific) for 10 minutes, prepared by diluting 16% paraformaldehyde in a cytoskeleton buffer consisting of 10 mM 2-N-morpholino ethane sulfonic acid (MES, pH 6.1, Fisher Scientific), 3 m_M_ MgCl_2_ (Fisher Scientific), 138 m_M_ KCl (Fisher Scientific), and 2 m_M_ EGTA (Sigma Aldrich). Next, cells were permeabilized using a 0.5% Triton-X (Fisher Bioreagents) solution in the cytoskeletal buffer for 5 minutes. Unreacted aldehydes were then blocked with 100 m_M_ glycine (Fisher Scientific) for 15 minutes. Cells were washed three times with 1x Tris Buffered Saline (TBS) for 5 minutes each. Cells were stained with Alexa 488-phalloidin (Thermo Fisher Scientific) for 1 hour and DAPI (Sigma) for 15-30 minutes in 1x TBS containing 0.1% (v/v) Tween-20 (Fisher Bioreagent) and 2% (w/v) Bovine Serum Albumin (BSA, Sigma Life Science). For phospho-myosin and anti-paxillin staining, after initial phalloidin staining and washing with 1x TBS, cells were incubated with 1:50 dilution of phospho-myosin antibody (Cell Signaling, mouse, 3675) or a 1:200 dilution of anti-paxillin primary antibody (BD Biosciences, mouse, 610051) for 24 hours at 4 °C.

Samples were then washed with 1x TBS, followed by incubation with a 1:200 dilution of donkey anti-mouse IgG secondary antibody (Jackson ImmunoResearch, 715-175-150) and 1:500 dilution of DAPI for 1 hour. Samples were washed again with 1x TBS three times for 5 minutes each. To improve image resolution, Prolong Gold mounting media (Thermo Fisher) was applied between the substrate with cells and the coverslips. Samples were sealed with VALAP (a mixture of Vaseline, lanolin, and paraffin) and imaged the next day using epifluorescence microscopy with 10x, 40x, and 60x objectives (*NA* = 0.3, *NA* = 1.3, and *NA* = 1.49, Nikon). The immunofluorescence images were analyzed with ImageJ. Lines around the cell’s edge were drawn and an oval was fit to the cell shape. The aspect ratio was calculated as the ratio of the major axis of the cell divided by the minor axis of the cell. The angle of the cell was calculated as the angle between the long axis of the cells and the horizontal vector of the image. To quantify the directionality of fixed cells, the migration vector was analyzed using Python. The Directionality Index (DI) was calculated as cos(2|Δ*θ*|), where ∣Δ*θ*∣ represents the angular deviation of each cell from the average cell orientation angle within the image. For cells with an area less than 300 µm² and an aspect ratio below 1.5, typically small, rounded cells whose direction could not be reliably determined, a random angle was assigned solely for the purpose of computing the average orientation angle. These cells were then individually assigned a DI value of 0 to reflect their lack of directional alignment.

### 5.8 Reverse Transcription Quantitative PCR (RT-qPCR)

We used a Luna Universal one-step RT-qPCR kit (E3005S, New England BioLabs, Ipswich, MA). According to the manufacturer’s protocol, 0.8 μg of total RNA was used in each reaction along with the SYBR green master mix, Luna WarmStart RT and 0.4 μM of each primer. The RT-qPCR reaction was programmed as follows: Reverse transcription cycle step was at 55 °C for 10 minutes, the initial denaturation step was at 95 °C for 1 minute, the subsequent denaturation steps were at 95 °C for 10 seconds, the extension step was 60 °C for 30 seconds and the melting curve step temperature was 95 °C for total of 40 cycles using the Oneplus Real-Time PCR system. Glyceraldehyde 3-Phosphate Dehydrogenase (GAPDH) was used as an internal control for normalization. GAPDH (490 bp): Forward (5’-GGGCTCTCCAGAACATCATC-3’) , Reverse (5’GACTGAGTGTGGCAGGGACT-3’); ITGα1 (333 bp): Forward (5’ AGACGTGCTGTTCATCTCGG 3’), Reverse (5’CACCAAGGCCCCCAATAGTC3’); ITGα2 (128 bp): Forward (5’TCAACCTCAAAATTCCTCTCCTGT 3’), Reverse (5’ ACCAACATCTTCAAAACTGTGCA 3’); ITGα10 (107 bp): Forward (5’ GAGAGCCCTGTACATGAGTGG 3’), Reverse (5’ GCAGTCTTTGTTTCTCGTCCC 3’); ITGα11 (104 bp): Forward (5’ GGAACGCTAAAGGATTCACACA 3’) , Reverse(5’ CCCACCACCACGTCATTGTA 3’); ITGβ1 (208 bp): Forward (5’ TGTGTTCAGTGCAGAGCCTT 3’), Reverse (5’ CCATGACCTCGTTGTTCCCA 3’); Discoidin Domain Receptor 1 (DDR1) (136 bp): Forward (5’ CTGTCTTACACCGCCCCTG 3’), Reverse (5’ CCACCACACCATCTGCCAG 3’) and DDR2 (122 bp), Forward (5’ GCAGATTGACTTGCACACCC 3’), Reverse (5’ GAGTGCCATCCCGACTGTAA 3’). Relative expression of the target genes was calculated by dividing 2^-*Ct*^ of the target gene with 2^-*Ct*^ of GAPDH, where *C_t_* is the threshold cycle number.

### 5.9 Western blotting

We employed Western blotting to quantify phosphorylated myosin expression across different cell lines. Using 2 x 10^6^ cells from different mammalian cell lines (HFF, HaCaT, MDA-MB-231, WM-266-4, and THP-1 cells) at an approximate confluency of 80% in a cell culture dish, cells are washed with 1x PBS and trypsinized, counted and centrifuged in a conical tube to form a pellet. A total of 100 μl of 1x sample buffer was added to the tube. The cells in the tube were lysed on ice with a conical pestle to ensure cells formed a uniform lysate. Protein lysate was heated at 94 °C for 5 minutes. Proteins were separated on 4-20% precast polyacrylamide gel (Bio-Rad, Min-PROTEAN® TGX Stain-Free™ Protein Gels, 10 well, 30 µl, 4568093) by loading a total of approximately 90 μg of protein lysate for each cell line in the gel wells, running the gel at 120 V for 1 hour and transferring it onto a polyvinylidene difluoride (PVDF) transfer membrane (Thermo-Fisher Scientific, PB9220) at 100 V for 1 hour and 30 minutes. The blot was blocked in PBST, and 5% skim dried nonfat milk at 37 °C for 1 hour. The membrane was incubated in primary phospho-myosin antibody (Cell Signaling, mouse, 3675, 1:200 dilution) and or β-actin antibody (Santa Cruz, mouse, C-2, sc-8432, 1:500 dilution). Antibodies were incubated at 4 °C overnight, washed using washing buffer three times for 5 minutes, followed by treatment with donkey anti-mouse secondary antibody (Jackson ImmunoResearch, 715-035-150, 1:500) incubated for 1 hour, then the membrane was rewashed three times for 5 minutes. Blots were developed using freshly prepared enhanced chemiluminescent (ECL) reagents (250 m_M_ luminol in DMSO, 90 m_M_ p-coumaric in DMSO acid,1 _M_ Tris-HCI pH 8.5, 30% H_2_O_2_ and water) and visualized using Bio-Rad Gel Doc XR+ Imaging system. To quantify the protein bands, we used Image J software. Band intensities on the immunoblot were quantified by selecting a rectangular region of interest around the band as well as one around a local background region. The intensity of the background region was subtracted from the intensity of the band region. This value was divided by the corresponding background subtracted value for β-actin as a loading control.

### 5.10 Quantifying gel compaction

For the gel compaction experiment, 10^6^ cells ml^-1^were embedded in a 2 mg ml^-1^ or 1.5 mg ml^-1^ collagen gel (500 µl) in DMEM and either 10% or 15% FBS depending on the specific cell line tested. To neutralize the acidic collagen gel compaction mixture, 0.1 _M_ filter sterile NaOH was employed. After thoroughly mixing the collagen gel compaction mixture and avoiding air bubble formation, the mixture was put into a pre-sterilized cylinder placed in a 6-well dish. The gel was incubated for 2 hours at 37 °C for gel formation, the cylinder was removed and 2 ml of cell culture media was added to the gel to produce a floating gel in media. Gel images were taken at 0, 4, 20, 32, 54, 72, 96, 132, and 140 hours to observe the change in gel diameter. For the SMIFH2 and CK-666 inhibitor-treated gel compaction experiments, final concentrations of 20 and 50 µ_M_ SMIFH2 and 5 and 25 µ_M_ CK-666 were achieved within both collagen gel and media around the gel. Images were analyzed in ImageJ to measure the gel diameter at different times. The gel diameter at different time points was analyzed using GraphPad Prism 8, where non-linear regression analysis for a two-phase decay model was performed. To determine the fractional compaction, the gel diameter at 48 hours for each replicate was calculated using the two-phase decay equation. The fractional compaction is defined as the ratio of the change in diameter from 0 hours to 48 hours, relative to the diameter at 0 hours.

### 5.11 Live cell migration imaging and analyses

Chambers were fabricated by cutting adhesive thin transparent silicone rubber sheets using a craft cutter to form a chamber on a glass surface. This chamber is filled with clear media, supplemented with 10-15% FBS, depending on the cell line. Once the media is prepared, the substrate with the cells was flipped over onto the chamber. The chamber was sealed with VALAP. The setup was then imaged over an 8-hour period with images captured at 2-minute intervals. This allowed for detailed observation and analysis of cell migration behavior. The DI calculations for live cell migration were based on the average of cos(2|Δ*θ*|), with ∣Δ*θ*∣ being the angle difference between the collagen fibril and the cell migration direction, calculated from changes in *x* and *y* positions at 2-minute intervals and averaged over 8 hours.

Speed was calculated by averaging the displacements measured every 2 minutes, dividing the total displacement over the 2 minutes by the corresponding time interval, and then averaging the resulting speeds over 8 hours. To categorize migration direction during live cell migration, the direction of cell movement was calculated over a 12 minute window every six 6 minutes. The number of steps was classified based on deviation angles: less than 30°, between 30° and 60°, and greater than 60°, using R for analysis. Steps with speeds less than 10 µm hour^-1^ were considered slow.

### 5.12 Assessing statistical significance

Qualitative statistical significance was assessed using a 95% confidence interval, calculated from the experimental data and placed on graphs in the form of error bars. Nonoverlapping confidence intervals indicate statistical significance to approximately *p* < 0.05. Quantitative statistical significance was assessed using the Wilcoxon rank-sum test, which is necessary to compare groups for non-normally distributed data. A binomial test was used for testing statistical significance in data including fraction of cell steps. All analyses were performed in R. For data sets with fewer than 10 data points per group, a one-way ANOVA (single factor) test was performed to assess statistical differences between conditions. In general *p* > 0.05 is denoted by n.s., *p* ≤ 0.05 is denoted by *, *p* ≤ 0.05 is denoted by ** and *p* ≤ 0.001 is denoted by ***. However, there are times when denoting statistical significance for each pairwise comparison would be complicated, particularly in situations where all pairwise comparisons are statistically significant. Consequently, in those situations the figure caption indicates more specifics about statistical comparisons.

## Abbreviations

DPBS: Phosphate-Buffered Saline
BME: (β-MercaptoEthanol)
APS: Ammonium Persulfate
APTES: 3-AminoPropylTriEthoxySilane
EGTA: Ethylene Glycol bis(2-aminoethyl ether)-N,N,N′,N′-Tetraacetic Acid
PVDF: Polyvinylidene DiFluoride
BSA: Bovine Serum Albumin
ECM: Extracellular Matrix
TEMED: N,N,N’, N’-Tetraacetylethylenediamines
TBS: Tris Buffered Saline
HEPES: 2-hydroxyethylpiperazine-1-ethane sulfonic acid
FBS: Fetal Bovine Serum
DMSO: DiMethyl SulfOxide
MES: 2-N-Morpholino Ethane Sulfonic acid
ECL: Enhanced ChemiLuminescent
HFF: Human Foreskin Fibroblast
EMT: Epithelial to Mesenchymal Transition
AFM: Atomic Force Microscopy
p-myo: phosphorylated myosin
PAA: PolyAcrylAmide
DDR: Discoidin Domain Receptors
GAPDH: Glyceraldehyde 3-Phosphate Dehydrogenase
af: Aligned Fibrils
tf: Transferred Fibrils
mdf: MultiDirectional Fibrils rf: Random Fibrils
m: Mica
g: Glass
0.2: 200 Pa PAA gel
2: 2000 Pa PAA gel
20: 20,000 Pa PAA gel

## Supporting Information

Supporting Information is available from the Wiley Online Library or from the author.

## Author Contributions

A.F.: Conceived experiments, conducted experiments, analyzed the data and interpreted the data across all figures, prepared the figures, wrote the article and edited the article. S.T.: Wrote code and analyzed the data in **Figures 1-8** and **Supplementary Figures 2-6**. S.A.S.: Conceived experiments, conducted experiments, analyzed the data and interpreted the data in **Figures 3-5** and **Supplementary Figures 3, 5&6**, wrote article and edited the article. F.R.N.: Conceived experiments, conducted experiments, analyzed data and interpreted the data in **Figure 3** and **Supplementary Figure 3&4**, wrote article and edited the article. M.F.: Conceived experiments, conducted experiments, analyzed the data and interpreted the data in **Supplemental Figure 1**, wrote the article and edited the article. E.C.: Conceived experiments and interpreted the data in **Supplemental Figure 1** and edited the article. I.C.S.: Conceived experiments, analyzed the data and interpreted the data across all figures, wrote the article, prepared the figures and edited the article.

## Acknowledgements

We acknowledge Curtis Mosher and the Roy J. Carver High Resolution Microscopy Facility and David Wright at the DNA facility. This work was supported by the National Institutes of Health under grant No. GM143302.

## Data Availability Statement

The data that support the findings of this study are available from the corresponding author upon reasonable request.

