## Supporting Data for "Formins and Arp2/3 Reciprocally Regulate Contact Guidance on Aligned Collagen Fibrils"

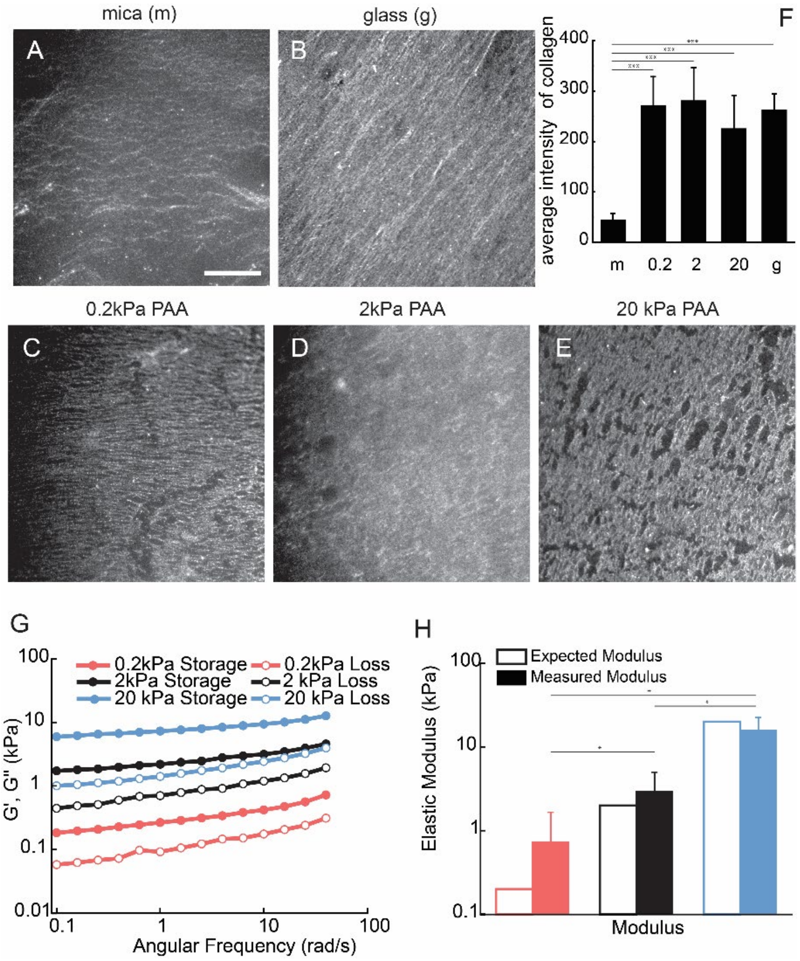


**Supplementary Figure 1: Collagen fibrils were uniformly transferred to all substrates**. (**A-E**) Immunofluorescence images of collagen fibrils on different substrates. (**F**) Quantified immunofluorescence intensity of collagen fibrils on different substrates. Statistical significance was assessed by comparing mica and other treatment conditions. (**G**) Frequency sweep of polyacrylamide (PAA) gels with varying cross-linker concentrations showing storage modulus (G′) and loss modulus (G″). (**H**) Quantification of elastic modulus calculated from storage modulus values across replicates for each gel condition. Statistical significance was assessed between all pairwise comparisons. Comparisons not shown were not statistically significant. Error bars represent 95% confidence intervals.

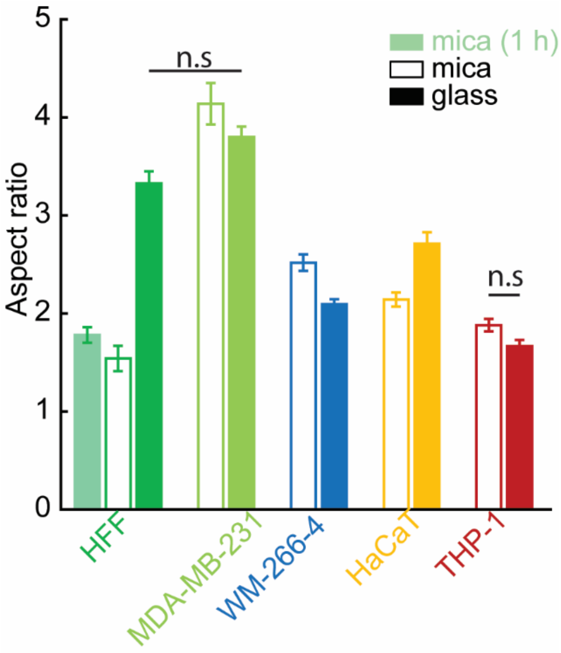


**Supplementary Figure 2: Cells exhibit varying elongation on mica and glass**. The aspect ratio of all cell lines on both mica (m) and glass (g). Statistical significance was assessed by comparing mica and glass conditions as well as each cell line. All comparisons showed statistically significant differences with $p \leq 0.001$, except for the comparisons explicitly marked on the plot. Error bars represent 95% confidence intervals.

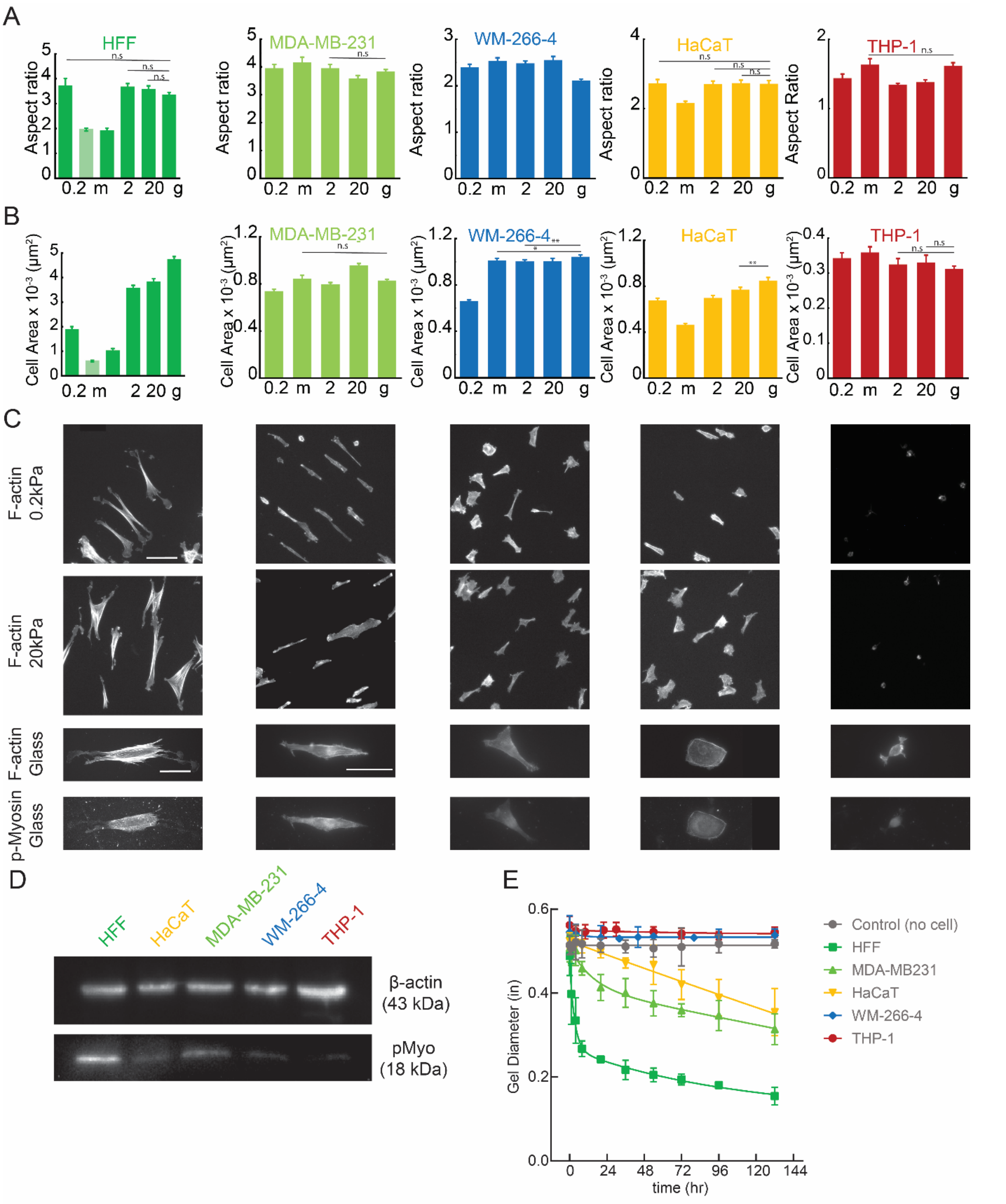


**Supplementary Figure 3: Different cells exhibit distinct morphologies on various substrates. (A)** aspect ratio and (**B**) cell area of all of the cell lines fixed on different substrates including 200 Pa PAA gel (0.2), mica (m), 2000 Pa PAA gel (2000), 20000 Pa PAA gel (20) and glass (g). Statistical significance was assessed by comparing all conditions to the glass condition. All comparisons showed statistically significant differences except for the ones explicitly shown on the plot. (**C**) Representative images the F-actin of all cell lines on 200 Pa and 20 kPa PAA gels and the high-resolution image of F-actin and phosphomyosin of all cell lines fixed on the glass. (**D**)Western blot example showing phosphomyosin and β-actin levels in all cell lines. (**E**) Gel diameter of compaction assay as a function of time for all of the cell lines. Error bars represent 95% confidence intervals.

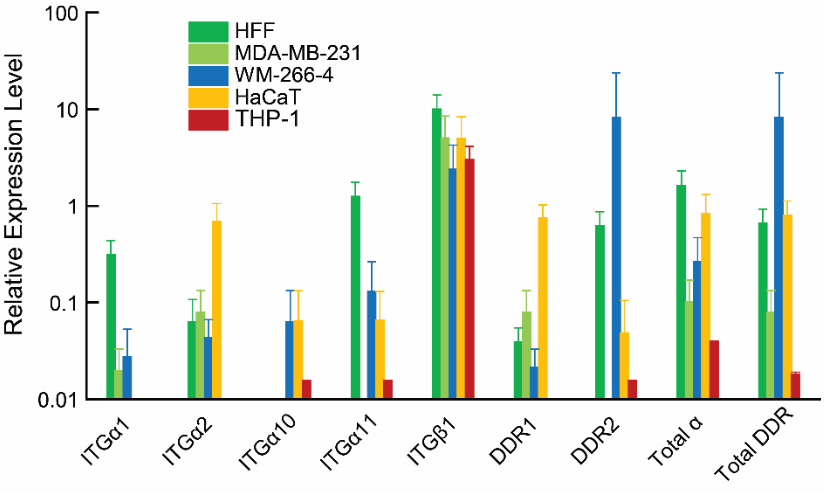


**Supplementary Figure 4:** Comparative qRT-PCR analysis of integrin and DDR receptor expression across five cell lines. mRNA levels of integrin α1, α2, α10, α11, β1, DDR1, DDR2, total α-subunits and total DDR relative to GAPDH were measured in human foreskin fibroblasts (HFF, green), MDA-MB-231 (light green), WM-266-4 (blue), HaCaT (yellow) and THP-1 (red) using qRT-PCR. Data not shown had expression levels lower than 0.01. $N_{samples} = 4$. Error bars represent the 95% confidence interval.

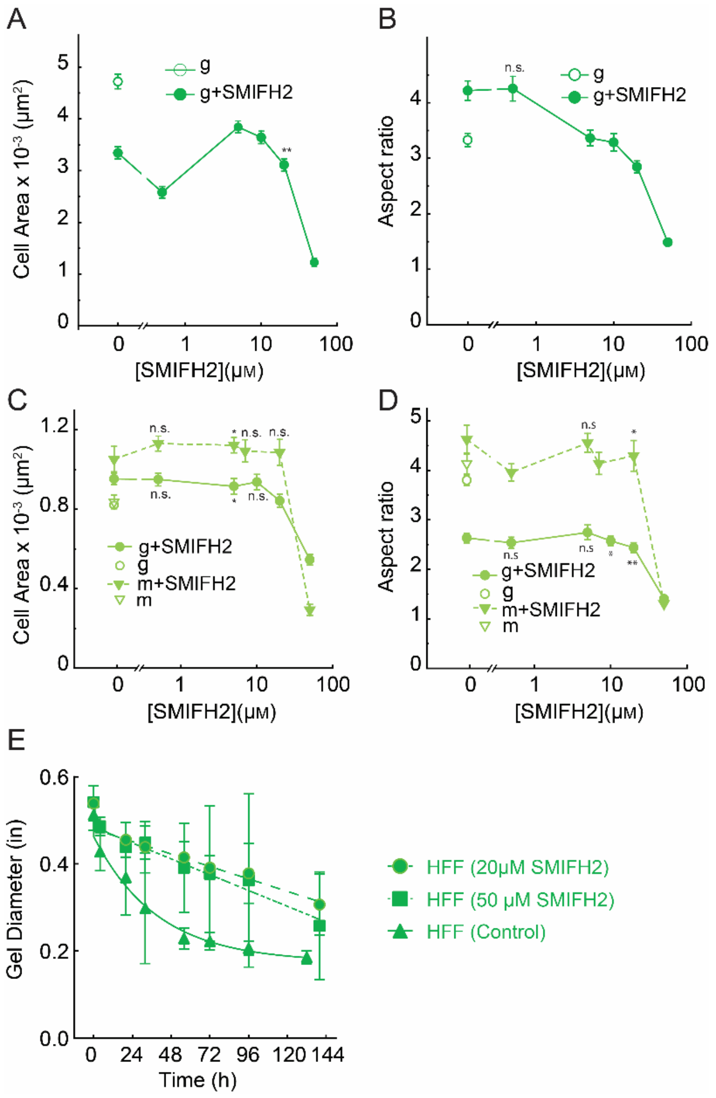


**Supplementary Figure 5: Formin inhibition alters cell morphology to a more rounded shape with a smaller area.** (**A**) Cell area and (**B**) aspect ratio of HFFs as a function of SMIFH2 concentration on glass. (**C**) Cell area and (**D**) aspect ratio of MDA-MB-231 cells as a function of SMIFH2 concentration on mica and glass substrates. Statistical significance was assessed by comparing 0.1% DMSO (control) and other treatment conditions. Comparisons not shown were statistically significant ($p \leq 0.001$). (**E**) Gel diameter as a function of time for all cell lines. Error bars represent 95% confidence intervals.

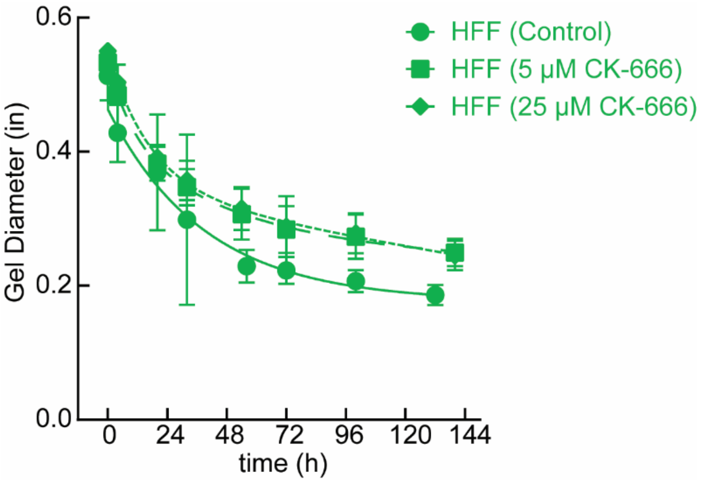


**Supplementary Figure 6: Arp2/3 inhibition results in reduced contractile force**. The gel diameter as a function of time for the HFF cells under both without drug condition and 5 μM CK-666. Error bars represent 95% confidence intervals.
